# A Protective Inter-Organ Communication Response Against Life-Threatening Malarial Anemia

**DOI:** 10.1101/2022.01.12.475857

**Authors:** Qian Wu, Euclides Sacomboio, Lara Valente de Souza, Rui Martins, Sílvia Cardoso, Temitope W. Ademolue, Tiago Paixão, Jaakko Lehtimäki, Caren Norden, Pierre-Louis Tharaux, Guenter Weiss, Fudi Wang, Susana Ramos, Miguel P. Soares

## Abstract

Anemia is a clinical hallmark and independent risk factor of malaria mortality, the disease caused by *Plasmodium spp.* infection. While malarial anemia arises from parasite-induced hemolysis, whether and how host metabolic adaptation to malaria regulates anemia severity is less understood. Here we demonstrate that reprogramming of organismal iron (Fe) metabolism by the kidneys is a central component of the host metabolic response regulating the pathogenesis of life-threatening malarial anemia. Renal proximal tubule epithelial cells (RPTEC) are the main cell compartment responsible for Fe storage and recycling during *Plasmodium* infection in mice. Transcriptional reprogramming of RPTEC couples immune resistance to *Plasmodium* infection to renal Fe export via the induction of the cellular Fe exporter SLC40A1/ferroportin 1. This integrated defense strategy is essential to deliver Fe to erythroblasts and support compensatory erythropoiesis to prevent the development of life-threatening anemia. Failure to mobilize Fe from RPTEC causes acute kidney injury (AKI) and is associated with life-threatening anemia in *P. falciparum*-infected individuals. These findings reveal an unexpected role of the kidneys in the control of organismal Fe metabolism and anemia severity during malaria.

## Introduction

Malaria is a vector-borne disease transmitted by the bite of a female *Anopheles* mosquito and characterized by the invasion of host red blood cells (RBC) by protozoan parasites of the *Plasmodium* genus. *Plasmodium* proliferation in RBC leads inexorably to hemolysis and to the development of more or less severe anemia, a clinical hallmark and independent risk factor of malaria mortality^1–3^.

Intravascular hemolysis produces extracellular hemoglobin (Hb), which releases its prosthetic heme groups upon oxidation^4, 5^. As it accumulates in plasma and urine during *Plasmodium* infection^6, 7^, labile heme acts as an alarmin^8^ and promotes the pathogenesis of severe malaria^4, 5, 9^.

The heme groups of Hb contain the largest proportion of bioavailable Fe in mammals and therefore, organismal Fe homeostasis relies on the recycling of this pool of Fe-heme^10^, by hemophagocytic macrophages in the red pulp of the spleen^11^. Once released from the protoporphyrin ring of heme, by the heme catabolizing enzyme heme oxygenase-1 (*HMOX1*/HO-1), Fe can be exported via *SLC40A1*^11^. Once exported from hemophagocytic macrophages Fe is bound by transferrin and delivered to erythroblasts in the bone marrow, supporting erythropoiesis and preventing the development of anemia^12^.

*Plasmodium* infection is associated with a transient depletion of hemophagocytic macrophages in the spleen^13^ and reprogramming organismal Fe metabolism^14^. This metabolic response relies on the induction of HO-1 in renal proximal tubule epithelial cells (RPTEC), where Fe is extracted from heme and stored by ferritin^7^. While essential to survive *Plasmodium* infection, this defense strategy does not exert a negative impact on the parasite^7^, establishing disease tolerance to malaria^15, 16^.

Here we describe that transcriptional reprogramming of RPTEC during malaria involves the induction of SLC40A1, which delivers the Fe stored in RPTEC to erythroblasts in the spleen. This is essential to support compensatory erythropoiesis and prevent the development of life-threatening anemia to establish disease tolerance to malaria. Failure to coordinate immune-driven resistance to *Plasmodium* infection with renal Fe mobilization leads to acute kidney injury (AKI) and is associated with life-threatening in *P. falciparum* infected individuals.

### Malaria is associated with Fe storage in RPTEC (*Fig. 1; S1-3*)

*Plasmodium chabaudi chabaudi* AS (*Pcc* AS) infection was associated with Fe accumulation in the kidneys of C57BL/6J mice (*Fig. 1A; S1A,B*), specifically at RPTEC luminal surface (*Fig. 1B,C; S1A*). Concomitantly, splenic Fe content was reduced (*Fig. 1A; S1C*), while hepatic (*Fig. 1A; S1D*) and cardiac (*Fig. 1A; S1E*) Fe content were marginally increased. This suggests that RPTEC are the main cell compartment responsible for Fe storage during *Plasmodium* infection.

**Figure 1.**
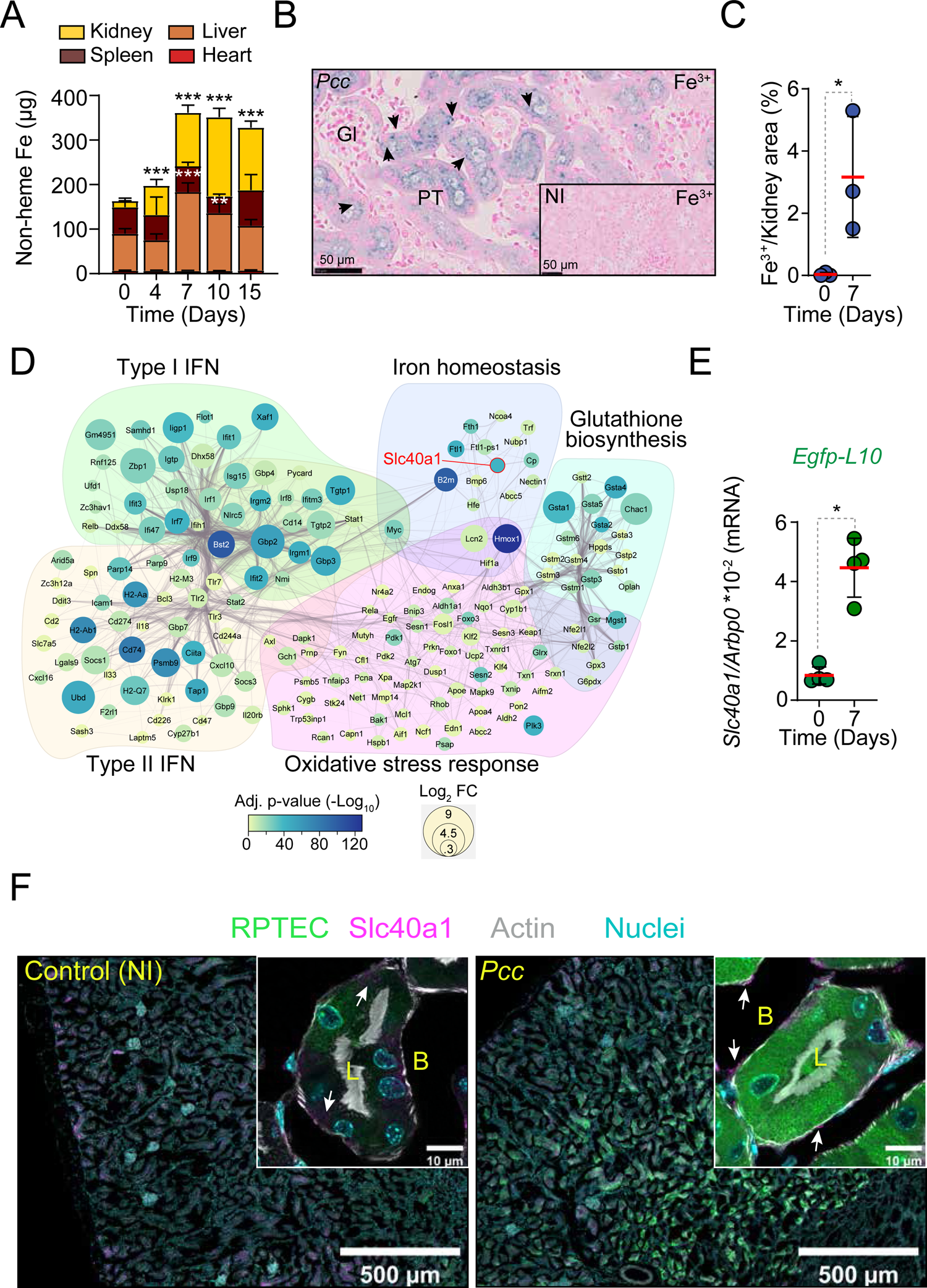
Malaria is associated with Fe storage in RPTEC. **A)** Total non-heme Fe content (mean µg ± SD) in organs from male C57BL/6 mice, before (D0) and after *Pcc* infection (N=4-5 mice *per* group). Data from one experiment. **B-C)** Perl’s Prussian Blue stain of non-heme Fe in kidneys from C57BL/6 mice, 7 days after *Pcc* infection, representative of 3 mice in 1 experiment. The indented rectangle corresponds to a non-infected (NI) C57BL/6 mouse. Arrowheads indicate Fe^3+^ (blue). Gl: Glomerulus, PT: proximal tubules. Scale bars: 50µm. **C)** Quantification of non-heme Fe staining *per* area of kidney section shown as mean (red bar) ± SD (N=3 *per* group). Circles correspond to individual mice. **D)** STRING database network analysis of genes belonging to the gene ontology/functional terms represented, which are upregulated upon *Pcc* infection. Edge thickness represents interaction confidence as reported by the STRING database v11.5. **E)** Expression of *Slc40a1* normalized to *Arbp0* ribosomal-associated mRNA in RPTEC of non-infected (Day 0) and *Pcc*-infected (Day 7) *Egfp-L10^Pepck^* mice quantified by qRT-PCR. Data represented as mean ± SD, in a subset of mice (N=4 group) from the same experiment as (D). Circles correspond to individual mice. **F)** Immunofluorescence imaging of kidney sections from control (NI, non-infected) *vs. Pcc*-infected (day 7) *Egfp-L10^Pepck^* mice, expressing eGFP specifically in RPTEC (green). Slc40a1 (magenta) was stained using a polyclonal antibody, DNA (nuclei) with DAPI (cyan) and actin (grey) was labeled using fluorescent-conjugated Phalloidin. White arrowheads indicate Slc40a1 expression. Images are representative of 3 animals *per* group from 2 independent experiments. (B) indicates basolateral and (L) luminal RPTEC surfaces. P values in (A) were determined using one-way ANOVA and in (C) and (E) using Mann Whitney test. *: P<0.05; **: P<0.01; ***; P<0.001.

To address how RPTEC increase their Fe storage capacity we analyzed the transcriptional response of RPTEC during *Pcc* infection in *Egfp-L10^Pepck^* mice expressing an EGFP-tagged L10 ribosomal subunit specifically in RPTEC (*Fig. S2A*)^17^. Analysis of mRNAseq data revealed that, at the peak of *Pcc* infection, there were 1717 genes repressed and 2261 induced in RPTEC (adjusted p-value <0.05; *Fig. S2B, Table S1*). The induced genes included type I and II interferon-responsive transcription factors (*e.g., Irf1, 2, 3, 7, 8 and 9*), cell-autonomous defense response genes (*e.g., Gbp*, 2, 3, 10 *Gsta1*, 2, 5) as well as genes involved in antigen presentation by major histocompatibility class I (MHC I; *e.g., H2K1*) and MHC II (*e.g., H2-DMb2, CD74*)(*Fig. 1D; S2C, Table S2*). An additional transcriptional program regulated by the nuclear factor kappa B (NF-κB) and associated with cellular responses to pathogen recognition and cytokines was also observed (*Fig. S2D, Table S2*). Moreover, there was a prominent oxidative stress-response (*Fig. 1D; S2D, Table S2*), orchestrated by the transcription factor erythroid 2–related factor 2 (NRF2/NFE2L2) and associated with the induction of genes involved in glutathione metabolism, metal chaperoning (*e.g., Mt1, Mt2*), heme catabolism (*e.g., Hmox1*), Fe storage (*e.g., Fth* and *Ftl* subunits of the ferritin complex) and cellular Fe export (*e.g., Slc40a1*) (*Fig. 1D; S2C, Table S2*). Conversely, mitochondrial genes were downregulated in RPTEC following *Pcc* infection (*Fig. S2E, Table S2*). This suggests that during *Plasmodium* infection RPTEC are reprogrammed to present antigens derived from pathogens^18^ while developing the ability to store and recycle catalytic Fe.

We hypothesized that Slc40a1 induction by RPTEC is required to mobilize the large amounts of Fe that accumulate in the kidneys during *Plasmodium* infection (*Fig. 1A-C*)^14^. We confirmed by qRT-PCR that the induction of renal *Slc40a1* mRNA in *Pcc*-infected *vs.* naïve C57BL/6J mice (*Fig. S3A*)^7^ occurs specifically in RPTEC, as determined in *Pcc*-infected *vs.* naïve *Egfp-L10^Pepck^* mice (*Fig. 1E*).

Labile heme was sufficient *per se* to induce the expression of *Slc40a1* mRNA (*Fig. S3B*) and protein (*Fig. S3C,D*) in primary mouse RPTEC *in vitro*. This suggests that labile heme contributes to reprogramming Fe metabolism in RPTEC to favor intracellular Fe export.

Induction of renal Slc40a1 protein during *Pcc* infection in C57BL/6J mice was confirmed by western blot (*Fig. S3E*) and associated with the RPTEC basolateral surface, as assessed by Slc40a1 and GFP (*i.e.,* RPTEC) co-immunostaining in *Egfp-L10^Pepck^* mice (*Fig. 1G, S4*).

### RPTEC *Slc40a1* is essential to establish disease tolerance to malaria (*Fig. 2; S5,6*)

To determine whether the RPTEC *Slc40a1* impacts the pathologic outcome of *Plasmodium* infection, we generated *Slc40a1^fl/fl^* mice carrying a functional knocked-in *Slc40a1^fl/fl^* allele (*Fig. S5A*). These were crossed with *Pepck^Cre/Wt^* mice^17^ (*Fig. S5A,B*) to generate *Pepck^Cre^Slc40a1^fl/fl^* mice, repressing the expression of the *Slc40a1^fl/fl^* allele in RPTEC from *Slc40a1^PepckΔ/Δ^* mice, upon exposure to acidified water (*Fig. S5C,D*)^17^. To confirm *Slc40a1^fl/fl^* deletion, *Pepck^Cre^Slc40a1^fl/fl^* mice were crossed with *Egfp-L10^fl/Wt^* mice^19^ to generate *Slc40a1^PepckΔ/Δ^Egfp-L10^Pepck^* mice, expressing an EGFP-tagged L10 ribosomal subunit specifically in RPTEC (*Fig. S5E*). Suppression of *Slc40a1* mRNA expression in RPTEC was confirmed by qRT-PCR of ribosomal-associated mRNA from *Egfp-L10^Pepck^ vs. Slc40a1^PepckΔ/Δ^Egfp-L10^Pepck^* mice (*Fig. S5F*).

**Figure 2.**
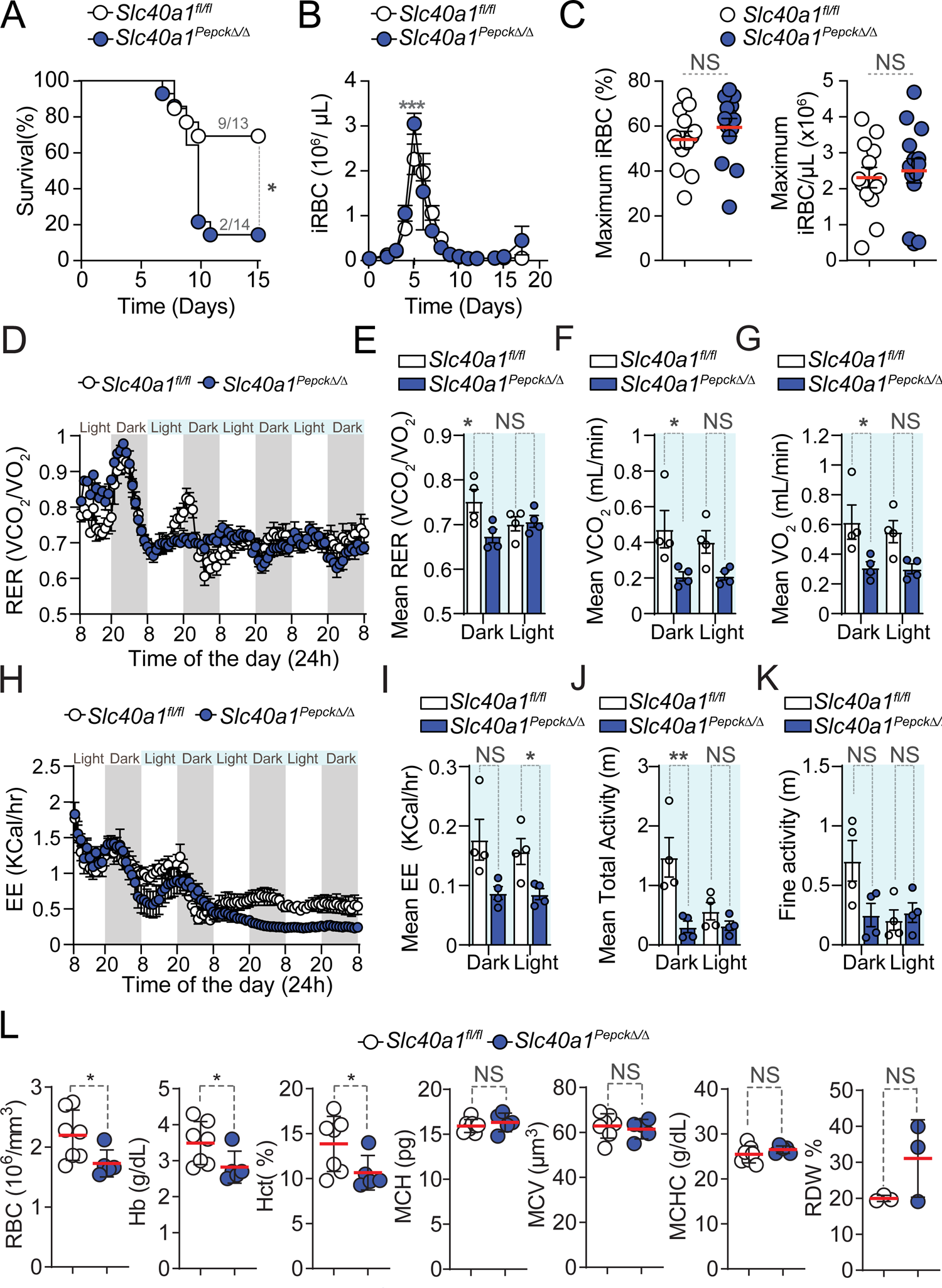
RPTEC Slc40a1 establishes disease tolerance to malaria. **A-C)** *Pcc* infection in *Slc40a1*^fl/fl^ (N=13) and *Slc40a1^Pepck^*^Δ/Δ^ (N=14) mice, pooled from four independent experiments with a similar trend. **A)** Survival. **B)** Pathogen load represented as mean number of iRBC per µL ± SEM. **C)** Maximum parasitemia (% iRBC) and pathogen load throughout the infection, represented as mean (red bar) ± SEM. Grey numbers in (A) indicate surviving over the total number of mice per genotype. **D-K)** Synchronized metabolic and behavioral quantification of *Slc40a1*^fl/fl^ (N=4, white circles) and *Slc40a1^Pepck^*^Δ/Δ^ (N=4, blue circles) mice, from 5-10 days after *Pcc* infection. Data from one experiment. **D)** Daily respiratory exchange ratio (RER) represented as mean ± SEM. **E-G)** Mean of daily averages of **E)** RER, **F)** VCO_2_ and **G)** VO_2_ in the period indicated in light blue in (D). Data is represented as mean ± SEM, segregated into daily light/dark cycle. **H)** Daily energy expenditure (EE), represented as mean ± SEM. **I-K)** Mean of daily averages of **I)** EE and **J)** total and **K)** fine activities in the period indicated in light blue in (D), represented as mean ± SEM, segregated into daily light/dark cycle. **L)** Hemograms from *Slc40a1*^fl/fl^ (N=7) mice and *Slc40a1^Pepck^*^Δ/Δ^ (N=5), 10 days post *Pcc* infection. Data represented as mean (red bar) ± SD, derived from 2 experiments. RBC: red blood cell count; Hb: hemoglobin concentration; Hct: hematocrit; MCH: mean corpuscular hemoglobin; MCH: mean corpuscular volume; MCHC: mean corpuscular hemoglobin concentration; RDW: RBC distribution width. Circles in (C), (E-G), (I-K) correspond to individual mice. P values of survival in (A) were determined using Log-rank (Mantel-Cox) test, in (B, D-K) using two-way ANOVA, and in (C, L) using Mann Whitney test. NS: not significant; *: P<0.05; **: P<0.01, ***:P<0.001.

Susceptibility to *Pcc* infection was increased upon *Slc40a1* deletion in RPTEC from *Slc40a1^PepckΔ/Δ^ vs. Slc40a1^fl/fl^* mice (*Fig. 2A*). This was not associated with changes in pathogen load (*Fig. 2B,C*), suggesting that RPTEC Slc40a1 contributes to the establishment of disease tolerance to malaria.

Deletion of *Slc40a1* exerted a negligible impact on transcriptional reprograming of RPTEC in response to *Pcc* infection (*Fig S6A, Table S3*), with 52 up-regulated and 77 down-regulated genes, compared to controls expressing *Slc40a1* in RPTEC (*Fig. S6A-C, Table S3*). The up-regulated genes were functionally related to histone acetylation and erythrocyte homeostasis, whereas down-regulated genes were associated with histone deacetylation (*Fig. S6C-D*). Principal component analysis showed that control *Egfp-L10^Pepck^* and *Slc40a1^PepckΔ/Δ^Egfp-L10^Pepck^* mice infected clustered independently of their genotype, while segregating based on *Pcc* infection (*Fig. S6E*). This suggests that the protective effect exerted by the induction of Slc40a1 in RPTEC is not exerted via a cell-autonomous mechanism that controls gene expression in RPTEC.

### *Slc40a1* expression by RPTEC is essential to support organismal O_2_/CO_2_ exchange and energy expenditure in response to malaria (*Fig. 2****; S7*)**

The lethal outcome of *Pcc* infection in *Slc40a1^PepckΔ/Δ^* mice was associated with a decrease of respiratory exchange ratio (RER; VCO_2_/VO_2_), which was more pronounced when compared to *Pcc*-infected *Slc40a1^fl/fl^* mice (*Fig. 2D, E*). This was driven by prominent decrease in the volumes of carbon dioxide (VCO_2_) emission (*Fig. 2F, Fig. S7A*) *vs.* oxygen (VO_2_) consumption (*Fig. 2G, S7B*) and was associated with lower energy expenditure (EE)(*Fig. 2H,I*) and total movement activity (*Fig. 2J*). While there was a tendency for reduction in fine movement activity (*Fig. 2K*) and lower food intake (*i.e.,* anorexia) (*Fig. S7C,D*), this was not statistically significant. Taken together these observations suggest that RPTEC Slc40a1 is necessary to sustain organismal RER and EE, above a threshold compatible with host survival. Of note, RPTEC Slc40a1 had no effect on steady-state food intake (*Fig. S7E,F*), total activity (*Fig. S7G*), RER (*Fig. S7H,I*), reflected by VCO_2_ (*Fig. S7J*) and VO_2_ (*Fig. S7K*), or EE (*Fig. S7L,M*) of *Slc40a1^PepckΔ/Δ^ vs. Slc40a1^fl/fl^* mice.

### RPTEC *Slc40a1* regulates malarial anemia (*Fig. 2, 3; S8-11*)

Based on the lower RER of *Pcc*-infected *Slc40a1^PepckΔ/Δ^ vs. Slc40a1^fl/fl^* mice, we hypothesized that beyond the possible effect of RPTEC Slc40a1 on organismal energy metabolism, the observed reduction in VO_2_ and CO_2_ exchange might also emerge from an exacerbation of anemia limiting gas exchange capacity. In support of this hypothesis, we found that at the peak (day 7, *Fig. S8A,B*) and the recovery phase (day 10, *Fig. 2L, S8B*) of *Pcc* infection, a more pronounced reduction of blood RBC count, Hb concentration and hematocrit in *Slc40a1^PepckΔ/Δ^ vs. Slc40a1^fl/fl^* mice was observed. This was not associated with changes in RBC parameters, including RBC mean corpuscular hemoglobin (MCH), mean cellular volume (MCV), mean cellular Hb content (MCHC) or red cell distribution width (RDW) (*Fig. 2L, S8A*).

**Figure 3.**
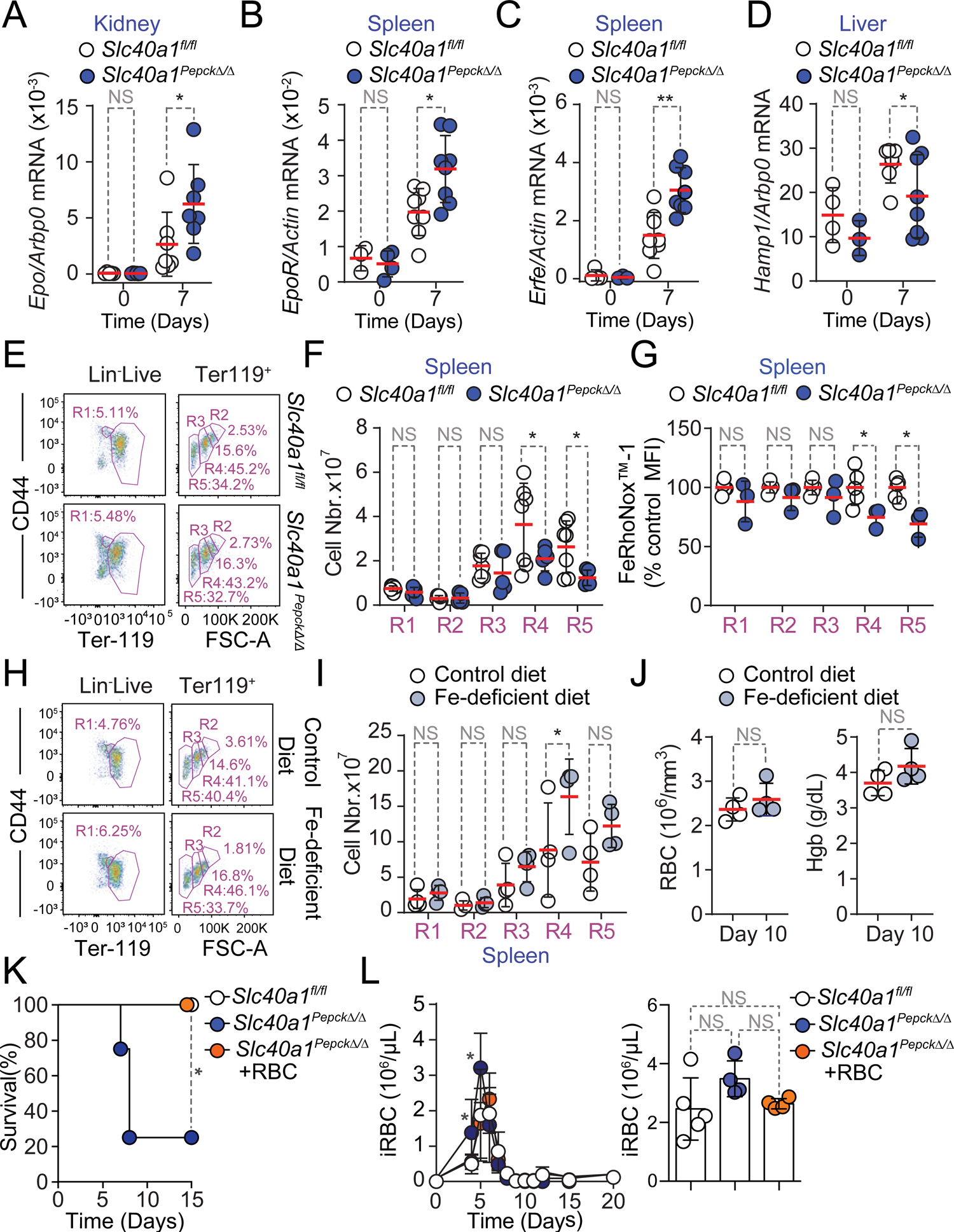
RPTEC Slc40a1 regulates the severity of malarial anemia. **A-D**) Quantification by qRT-PCR of mRNA encoding **A**) erythropoietin (*Epo*) in the kidney, **B**) Epo receptor (*EpoR*) in the spleen, **C**) Erythroferrone (*Erfe*) in the spleen and **D**) hepcidin (*Hamp1*) in the liver of *Slc40a1*^fl/fl^ mice not infected (Day 0; N=3-5) or infected with *Pcc* (Day 7; N=7-8) and *Slc40a1^Pepck^*^Δ/Δ^ mice not infected (Day 0; N=3-6) or infected with *Pcc* (Day 7; N=7-8). Data from 2 independent experiments. **E-F)** Splenic compensatory erythropoiesis in *Slc40a1*^fl/fl^ (N=5) and *Slc40a1^Pepck^*^Δ/Δ^ (N=7) mice, 10 days after *Pcc* infection. Data from 2 experiments. **E)** Representative FACS plots of the gating strategy based on size and CD44 and Terr119 expression used to identify different RBC developmental stages. Full gating strategy in *Figure S_1.* **F)** Quantification of cell numbers, represented as mean (red bar) ± SD, for the different cell populations identified in E). **G)** Relative quantification of Fe^2+^ (FeRhoNox^TM^-1 staining) in spleen (**Extended Data Figure 10A**, full gating strategy in *Figure S_3*), normalized to the average mean fluorescence intensity (MFI) of each erythroblast stage (*i.e.,* R1-5) in *Pcc*-infected *Slc40a1^fl/fl^* (N=5) and *Slc40a1^Pepck^*^Δ/Δ^ (N=3) mice, represented as mean (red bar) ± SD. Data from 2 experiments. **H-J)** Splenic compensatory erythropoiesis in C57BL/6 mice fed Fe-deficient (N=4) vs. control chow (N=4) diets, 10 days after *Pcc* infection. Data from one experiment. **H)** Representative FACS plots of the gating strategy based on size and CD44 and Terr119 expression used to identify different RBC developmental stages. Full gating strategy in *Figure S_1.* **I)** Quantification of cell numbers, represented as mean (red bar) ± SD, for the different cell populations identified in (H). **J)** Hemogram from the same mice as in (H). RBC: red blood cell count; Hb: hemoglobin concentration. **K-L)** Survival (**K**), pathogen load (**L,** left panel) and maximum pathogen load (**L,** right panel) during the course of *Pcc* infection in *Slc40a1^Pepck^*^Δ/Δ^ mice receiving or not (PBS; N=4) purified RBC (8×10^8^, i.p., N=4). *Pcc* infected *Slc40a1*^fl/fl^ (N=4) mice were used as controls. Data from 2 independent experiments. Data in A-D, F, H, I and K) represented as mean (red bar) ± SD. Circles in (A-D, F,G, I,J and L, right panel) represent individual mice. P values in (A-C, F, G, I), determined using one-way ANOVA, in (D) using Welch’s t test, in (J) using Mann Whitney test, in (K) determined with Log-rank (Mantel-Cox) test and in (L) using two-way ANOVA. NS: not significant; *: P<0.05; **: P<0.01.

The individual disease trajectories established by the relationship between circulating RBC numbers *vs.* pathogen load during *Pcc* infection (*Fig. S8C*), confirmed that *Slc40a1^PepckΔ/Δ^* mice reached lower numbers of circulating RBC, compared to *Slc40a1^fl/fl^* mice (*Fig. S8D*). This was associated with a more pronounced reduction of body temperature (*Fig. S8E-G*), without affecting maximum body weight loss (*Fig. S8H-J*). These observations are in line with clinical studies^20–23^, suggesting that *Plasmodium* is not the sole driver of malarial anemia severity and that Slc40a1 can regulate anemia severity without interfering with pathogen load^24^.

We next investigated how RPTEC Slc40a1 reprograms organismal Fe metabolism to limit malarial anemia severity. We found that mRNA expression encoding the renal erythropoiesis-inducing hormone erythropoietin (*Epo*) was induced to a greater extent in *Pcc*-infected *Slc40a1^PepckΔ/Δ^ vs. Slc40a1^fl/fl^* mice, compared to genotype matched non-infected controls (*Fig. 3A*). This was also the case for erythropoietin receptor (*EpoR*)(*Fig. 3B*) and erythroferrone (*Erfe*)(*Fig. 3C*) mRNA expression in the spleen. Concomitantly, there was a reduction of hepcidin (*Hamp1*) mRNA expression in the liver (*Fig. 3D*). These observations led to the hypothesis that RPTEC Slc40a1 reduces malarial anemia severity via a mechanism that increases Fe re-utilization and delivery to erythroblasts^25–28^.

*Pcc* infection was associated with the induction of compensatory erythropoiesis, characterized by the development of polychromatic erythroblasts into orthochromatic erythrocytes/reticulocytes and mature RBCs in the spleen (*Fig. S9A, S_1*). Compensatory erythropoiesis was impaired in *Pcc*-infected *Slc40a1^PepckΔ/Δ^ vs. Slc40a1^fl/fl^* mice (*Fig. 3E,F, S_1*), without affecting medullary erythropoiesis (*Fig. S9B,C*). This suggests that RPTEC Slc40a1 is essential to support compensatory erythropoiesis during malaria, without affecting medullary erythropoiesis.

Whether, similarly to sterile intravascular hemolysis^29^, hemophagocytic macrophages control the pathogenesis and/or progression of malarial anemia is not clear^13, 30^. Refuting this idea, the number of hemophagocytic macrophages containing intracellular RBC was indistinguishable in the spleen *(Fig. S10A, S_2),* liver *(Fig. S10B)* and kidneys (*Fig. S10C*) of *Pcc*-infected *Slc40a1^PepckΔ/Δ^ vs. Slc40a1^fl/fl^* mice. This suggests that renal RPTEC, rather than hemophagocytic macrophages, are the main cell compartment regulating the severity of malarial anemia.

In further support that Fe mobilization via RPTEC Slc40a1 is essential to support compensatory erythropoiesis, the intracellular Fe content of splenic erythroblasts was lower in *Pcc*-infected *Slc40a1^PepckΔ/Δ^ vs. Slc40a1^fl/fl^* mice (*Fig. 3G, S11A, S_3*). This was not observed in bone marrow erythroblasts (*Fig. S11A, B*), suggesting that Fe mobilization via RPTEC Slc40a1 is essential to support compensatory erythropoiesis.

We also tested whether Fe uptake from diet regulates compensatory erythropoiesis and malarial anemia severity. This hypothesis was not supported by the reduction of food intake (*i.e.,* anorexia) observed at the peak of *Pcc* infection (*Fig. S7C, D*). Moreover exposure to a diet with reduced Fe content, from day 7 to 10 after *Pcc*-infection, failed to compromise erythropoiesis (*Fig. 3H,I, S_1*) and to limit the severity of anemia (*Fig. 3J, S9D*). Moreover, RBC transfusion protected *Slc40a1^PepckΔ/Δ^* mice from succumbing to *Pcc* infection (*Fig. 3K*), sustaining thermoregulation *(Fig. S11C-E)*, without affecting weight loss *(Fig. S11F-H*) or pathogen load (*Fig. 3L*). Taken together, these observations provide further evidence for the critical role of Fe mobilization via RPTEC Slc40a1 in supporting compensatory erythropoiesis and limiting the development of life-threatening malarial anemia.

### RPTEC Fe export via *Slc40a1* suppresses acute kidney injury (AKI)(*Fig. 4; S12-15*)

Fe accumulation in the kidneys was increased in a sustained manner during *Pcc* infection in *Slc40a1^PepckΔ/Δ^ vs. Slc40a1^fl/fl^* mice (*Fig. 4A*), while hepatic Fe content was only marginally increased *Fig. S12A*), splenic Fe content was not effected (*Fig. S12B*) and cardiac Fe content was marginally reduced (*Fig. S12C*). Moreover *Pcc*-infected *Slc40a1^PepckΔ/Δ^* mice also had a more pronounced increase in Fe concentration in plasma (*Fig. 4B*), while transferrin saturation remained similar to *Pcc*-infected *Slc40a1^fl/fl^* mice (*Fig. S12D*). This suggests that a significant proportion of circulating Fe in *Pcc*-infected *Slc40a1^PepckΔ/Δ^* mice was not bound to transferrin (non transferrin bound Fe; NTBI) (*Fig. 4C*). In keeping with this observation *Pcc*-infected *Slc40a1^PepckΔ/Δ^* mice had lower levels of circulating transferrin, but not albumin, compared to *Pcc*-infected *Slc40a1^fl/fl^* mice (*Fig. 4D*), consistent with transferrin being a negative acute phase protein^31^. As NTBI fails to support erythropoiesis^32^, circulating Fe in *Pcc*-infected *Slc40a1^PepckΔ/Δ^ vs. Slc40a1^fl/fl^* mice should fail to counter the severity of malarial anemia.

**Figure 4.**
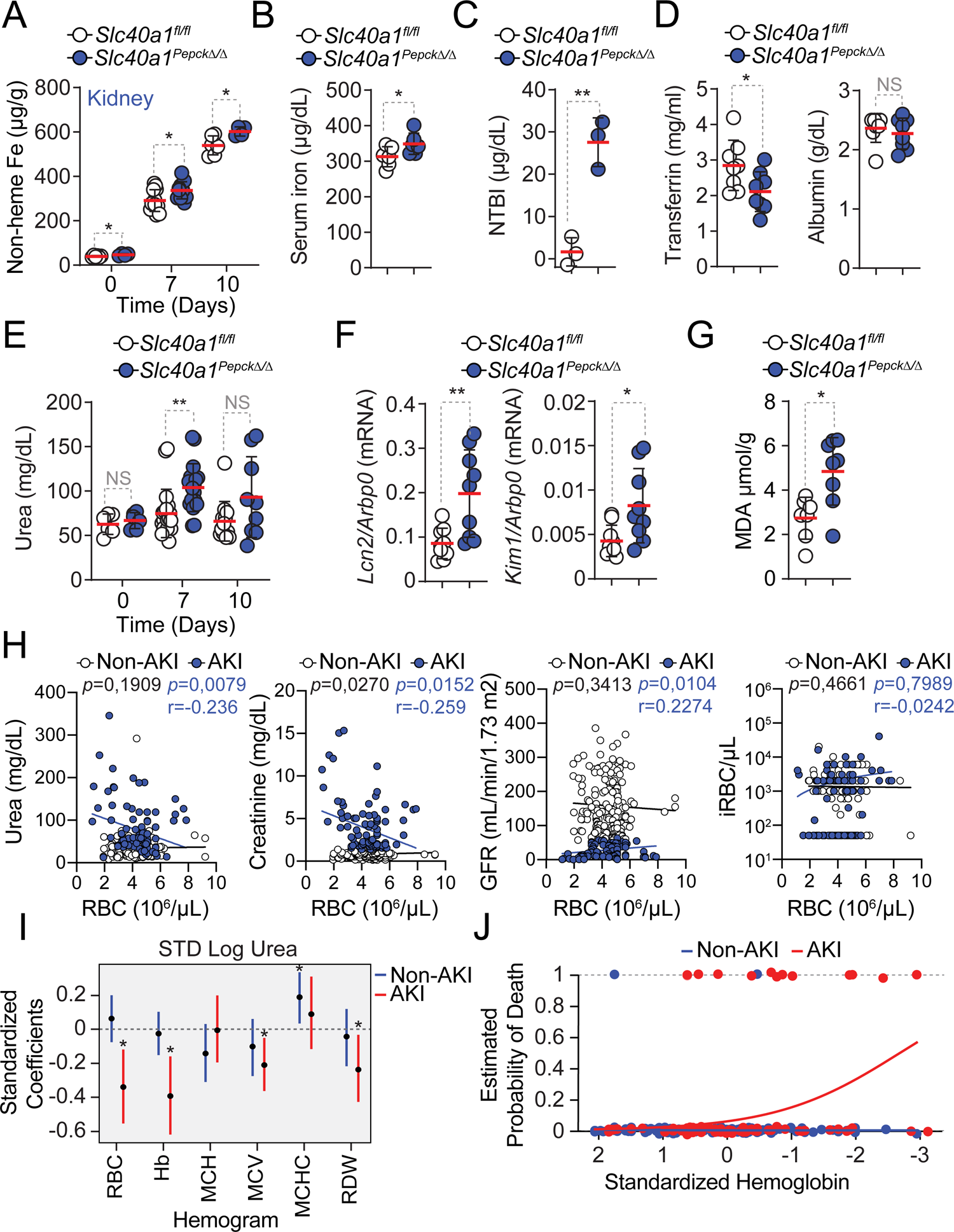
AKI predicts anemia severity in *P.* falciparum-infected individuals. **A)** Non-heme Fe content, represented as µg of Fe *per* g of tissue in the kidneys of *Slc40a1*^fl/fl^ mice and *Slc40a1^Pepck^*^Δ/Δ^, before infection (D0, N=6 vs N=4) or after *Pcc* infection (Day 7, N=12 *vs.* N=11; and Day 10, N=6 vs N=4). Data from 2 independent experiments with similar trend. Serum **B)** Fe in *Pcc*-infected (day 7) *Slc40a1*^fl/fl^ (N=6) and *Slc40a1^Pepck^*^Δ/Δ^ (N=6) mice, **B)** non-transferrin-bound iron (NTBI) in *Pcc*-infected (day 7) *Slc40a1*^fl/fl^ (N=3) and *Slc40a1^Pepck^*^Δ/Δ^ (N=3) mice and **D)** transferrin and albumin concentration in *Pcc*-infected (day 7) *Slc40a1*^fl/fl^ (N=8) and *Slc40a1^Pepck^*^Δ/Δ^ (N=8) mice. Data from 2 independent experiments. **E)** Serum urea concentration in the same mice as (A). **F)** Quantification by qRT-PCR of *Lcn2* and *Kim1* mRNA in the kidneys of *Pcc*-infected *Slc40a1*^fl/fl^ (N=9) and *Slc40a1^Pepck^*^Δ/Δ^ (N=9) mice, 7 days post-infection. Data from one experiment. **G)** Quantification of the lipid peroxidation product malondialdehyde (MDA) in the kidneys of *Pcc*-infected *Slc40a1*^fl/fl^ (N=7) and *Slc40a1^Pepck^*^Δ/Δ^ (N=8) mice, 7 days post-infection. Data from 2 independent experiments. **H)** Linear correlations between serum urea, creatinine, glomerular filtration rate (GFR) and parasitemia (iRBC/µL) *vs.* number of circulating RBC in *P. falciparum*-infected patients diagnosed with AKI or not (non-AKI) with corresponding *p* values (non-AKI: black; AKI: blue), and the Spearman’s correlation coefficient (r) for AKI patients. **(I)** Quantification of associations between urea and hemogram outputs after adjustment for gender, age and AKI status. Each bar depicts the estimate and uncertainty of the standardized coefficient (95% HPD) for urea of linear regression with the indicated hemogram output as response variable and urea, parasitemia, AKI status, age and sex as independent variables. **(J)** Probability of death as a function of anemia in AKI. Dots represent the actual data of death *vs*. standardized Hemoglobin concentration. Lines represent the logistic regression for the probability of death as a function of Hemoglobin concentration, stratified by AKI/non-AKI (red and blue colors, respectively), holding all other values constant at the mean of the population. Dashed line at Y=1 indicates death. Data in (A-G) represented as mean (red bar) ± SD. Circles represent individual mice. Circle in (H) represent individual patients. P values in (A,E) were determined using one-way ANOVA, in (B-D,F,G) using Mann Whitney test and in (H) using a Spearman’s rank correlation coefficient test. NS: not significant; *: P<0.05; **: P<0.01.

We then asked whether renal Fe export via RPTEC Slc40a1 regulates the pathogenesis of AKI, a common presentation and independent risk factor of malaria mortality^33–36^. This hypothesis was strongly supported by the higher accumulation of urea in the plasma of *Pcc*-infected *Slc40a1^PepckΔ/Δ^ vs. Slc40a1^fl/fl^* mice (*Fig. 4E*). This was not associated however, with an increase in creatinine concentration in plasma (*Fig. S13A*). On the other hand there was an increase in renal Lipocalin 2 (*Lcn2*) and kidney injury molecule 1 (*Kim1*) mRNA expression (*Fig. 4F*). Moreover, there was an accumulation of polyunsaturated fatty acid peroxidation (*i.e.,* malondialdehyde; MDA) (*Fig. 4G*), also associated with the pathogenesis of AKI^37^. This was not observed in the spleen (*Fig. S13B*).

*Pcc*-infected *Slc40a1^PepckΔ/Δ^* mice accumulated, albeit in a transient manner, alanine aminotransferase (ALT) (*Fig. S13C*) and aspartate aminotransferase (AST) (*Fig. S13D*) in plasma. This was also the case for lactate dehydrogenase (LDH) (*Fig. S13E*) but not Troponin 1 (*Fig. S13F*). This was not associated with overt hepatic (*Fig. S14A*), renal (*Fig. S14B*) or cardiac (*Fig. S14C*) histopathologic damage. Electron microscopy of the kidneys showed no evidence of epithelial ultrastructural alterations, with normal nuclei, cell polarity, endosomes, lysosomes and mitochondria density (*Fig. S15*). Of note, RPTEC from *Pcc*-infected *Slc40a1^PepckΔ/Δ^* mice presented an accumulation of electron-dese granules, consistent with consistent with lysosome accumulation and intracellular Fe deposits (*Fig. S15*). This suggests that Fe export from RPTEC via Slc40a1 limits intracellular Fe accumulation and counters the development of malarial AKI, likely impacting on the pathogenesis of life-threatening malarial anemia.

### AKI predicts anemia severity in *P. falciparum* infected individuals (*Fig. 4; S16*)

To address whether, similarly to that observed in experimental rodent malaria, malarial anemia severity in humans is also associated with renal dysfunction, we performed a retrospective analysis of clinical data from 400 individuals with confirmed *P. falciparum* infection. Among these 131 (32.7%) developed AKI and 269 (67.3%) did not (non-AKI) (*Table 1*). In line with KDIGO guidelines^38^, AKI was characterized by higher blood urea, blood urea nitrogen, blood creatinine concentration and lower glomerular filtration rate (GFR), than measured in non-AKI patients (*Tables 1, 2*).

**Table 1.** Baseline demographics and clinical characteristics of patients with severe falciparum malaria by AKI status at enrolment

| Variable | Total<br>(*n* = 400) | No AKI<br>(*n* = 269) | AKI<br>(*n* = 131) | *P* value |
| --- | --- | --- | --- | --- |
| **Demographics** |  |  |  |  |
| Age (years) | 25,0 (21-32) | 25,0 (21-32) | 25,0 (21-35) | 0,8513 |
| **Gender** |  |  |  |  |
| Female | 179 (44,8) | 115 (46,1) | 64 (42,0) | 0,2492<sup>b</sup> |
| Male | 221 (55,2) | 154 (53,9) | 67 (58,0) |  |
| **Blood group** |  |  |  |  |
| A<sup>a</sup> | 50 (24,9) | 24 (19,0) | 26 (34,7) | **0,0132<sup>b</sup>** |
| B<sup>a</sup> | 76 (37,8) | 52 (41,3) | 24 (32,0) | 0,1899<sup>b</sup> |
| AB<sup>a</sup> | 5 (2,5) | 4 (3,2) | 1 (1,3) | 0,4176<sup>b</sup> |
| O<sup>a</sup> | 70 (34,8) | 46 (36,5) | 24 (32,0) | 0,5165<sup>b</sup> |
| **Clinical parameters at enrolment** |  |  |  |  |
| Temperature (°C) | 37,00 (36-37,1) | 37,00 (36-37,08) | 37,00 (36-38,0) | 0,0637 |
| Weight (Kg) | 60,00 (56,78-70) | 60,0 (57-70) | 59,00 (56,7-70) | 0,6590 |
| Pulse rate (breaths/min) | 74,0 (63,75-86) | 75,0 (64,5-88) | 72,0 (62-82,25) | **0,0258** |
| Respiratory rate (breaths/min) | 22,0 (19,0-24,0) | 22,0 (19,0-23,0) | 22,0 (19,0-24,0) | 0,7332 |
| Oxygen Saturation (%) | 99,0 (99-99) | 99,0 (99-99) | 99,0 (99-99) | **0,0470** |
| **AKI stage at enrolment** |  |  |  |  |
| AKI 1<sup>a</sup> | 61 (15,25) | - | 61 (46,57) |  |
| AKI 2<sup>a</sup> | 15 (3,75) | - | 15 (11,45) |  |
| AKI 3<sup>a</sup> | 55 (13,75) | - | 55 (41,99) |  |

All values are **median** (IQR) unless otherwise specified, as in <sup>a</sup>: number (%). *P* values determined using Mann-Whitney U test or Chi-square test (<sup>b</sup>). Abbreviations: AKI: acute kidney injury. AKI diagnosis according to KDIGO guidelines, based on serum creatinine increase from baseline: AKI 1: 1,5-1,9 times baseline or $\geq 3,0$ mg/dL; AKI 2: 1,5-1,9 times baseline; AKI 3: 3 times baseline or $\geq 4,0$ mg/dL.

**Table 2.**
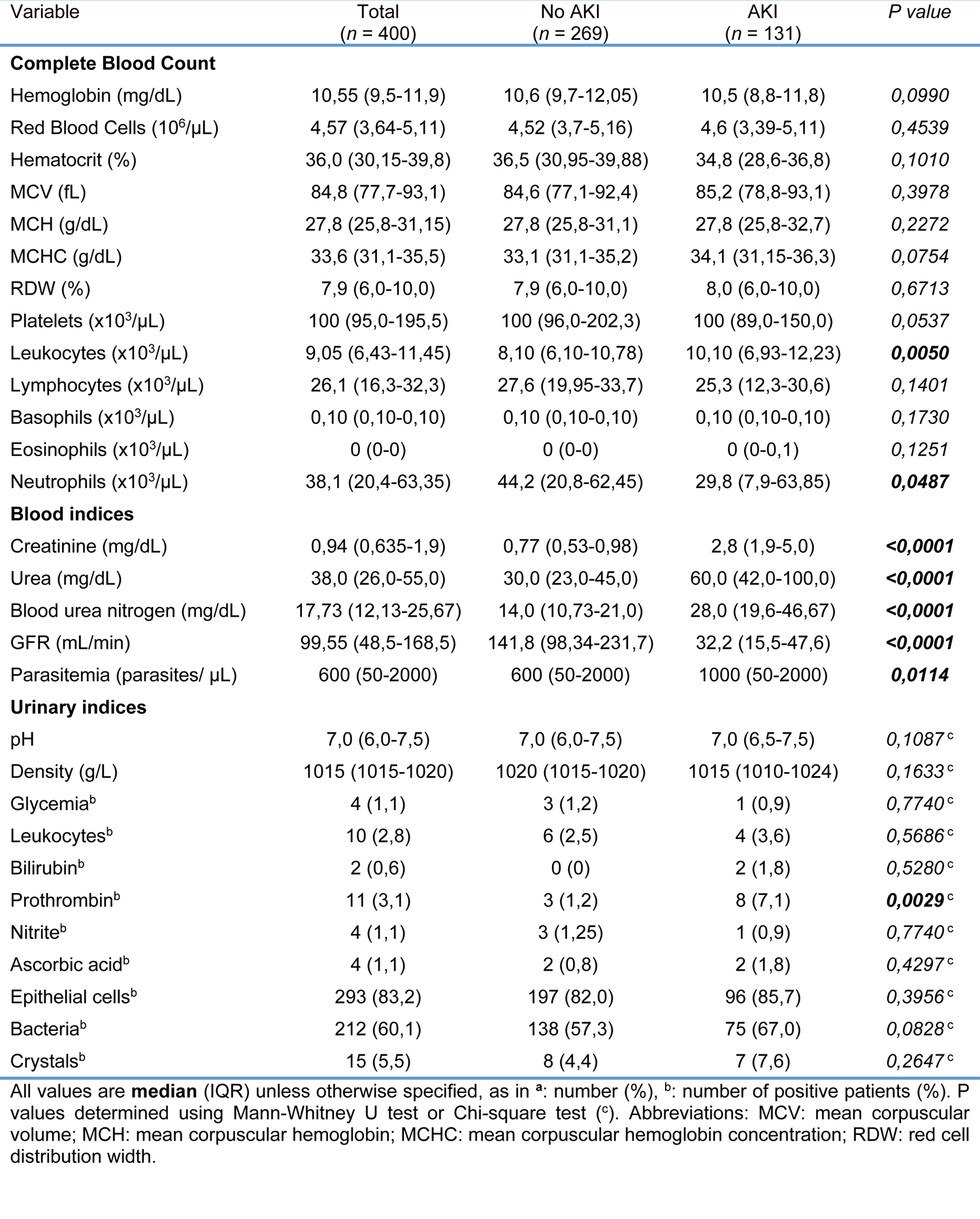
Baseline admission laboratory parameters of patients with severe falciparum malaria by AKI status at enrolment

AKI was associated with higher mortality rate (*Table 3*), consistent with previous clinical studies^36, 37^. Moreover, there was a significant negative correlation between blood urea concentration *vs.* blood RBC count (*Fig. 4H*), Hb concentration (*Fig. S16A*) and hematocrit (*Fig. S16B*) in AKI patients, not observed in non-AKI patients. A significant negative correlation between blood creatinine concentration *vs.* blood RBC count (*Fig. 4H*), Hb concentration (*Fig. S16A*) and hematocrit (*Fig. S16B*) was also observed in AKI but not in non-AKI patients. This was associated with significant positive correlation between GFR *vs.* blood RBC count (*Fig. 4H*), Hb concentration (*Fig. S16A*) and hematocrit (*Fig. S16B*) in AKI but not non-AKI patients. Blood urea concentration was a significant predictor of anemia severity in AKI but not in non-AKI patients, as shown by a negative standard coefficient in the regression analyses of blood RBC counts and Hb concentration (*Fig. 4H,I*). Taken together these observations argue for a direct association between AKI and the clinical severity of malarial anemia.

**Table 3.** Outcome by AKI status on enrolment

| Outcome | Total<br>(*n* = 400) | No AKI<br>(*n* = 269) | AKI<br>(*n* = 131) | *P* value |
| --- | --- | --- | --- | --- |
| Discharged (%) | 140 (35,0) | 98 (36,4) | 42 (32,1) | 0,3898 |
| Hospitalized (%) | 237 (59,25) | 169 (62,8) | 68 (51,9) | **0,0370** |
| Dead (%) | 23 (5,75) | 2 (0,7) | 21 (16,0) | **<0,0001** |

All values are number of patients (%). *P* values determined using Chi-square test. Abbreviations: AKI: acute kidney injury.

While AKI was associated with higher parasitemia (*Table 2*), there was no significant correlation between parasitemia *vs.* blood RBC count (*Fig. 4H*), Hb concentration (*Fig. S16A*) or hematocrit (*Fig. S16B*), among AKI patients. Moreover, parasitemia (*Fig. 4H, S16C*) was not a significant predictor of anemia severity, similar to gender (*Fig. S16D*) or age (*Fig. S16E*). This suggests that while AKI carries a significant impact on the severity of malarial anemia, this pathogenic effect is exerted irrespectively of parasitemia, consistent with previous observations^20–23^. Moreover, there was a significant association between probability of death and anemia (*i.e.* lower hemoglobin levels) in AKI but not in non-AKI patients (*Fig. 4J*). This suggests that AKI contributes to the development of anemia increasing the probability of death in *P. falciparum*-infected malaria patients.

## Discussion

It is well established that RPTEC express Fe-sensing and Fe-regulatory genes constitutively^39^. However, under steady-state conditions this does not appear to endow the kidneys with a major role in the regulation of organismal Fe metabolism. In sharp contrast, the kidneys become central to the regulation of organismal Fe metabolism during malaria, suppressing the development of life-threatening malarial anemia. This relies on the transcriptional reprogramming of RPTEC, partaking in “salvage pathway” to overcome the relative loss of heme catabolizing and Fe recycling capacity provided at steady state by hemophagocytic macrophages^13^. This unsuspected pathophysiologic role if RPTEC is essential to store and recycle Fe to suppress the development of life-threatening anemia while also preventing the pathogenesis of AKI^40^. We found that the later is a significant predictor of anemia severity and mortality in *P. falciparum*-infected patients. These findings are consistent with intravascular hemolysis partaking in the pathogenesis of severe falciparum AKI^37^ and provide a plausible explanation for previous associations between AKI and malarial anemia severity^35, 41^. The protective effect exerted by Fe recycling from RPTEC, via the induction of SLC40A1, acts irrespectively of parasite load, which might contribute to explain the recent association of the Q248H *SLC40A1* “gain-of-function” mutation with lower incidence of anemia in endemic areas of malaria, without interfering with *P. falciparum*^24^.

## Supporting information

western blot raw data

## Extended Data Figure

**Extended Data Figure 1.**
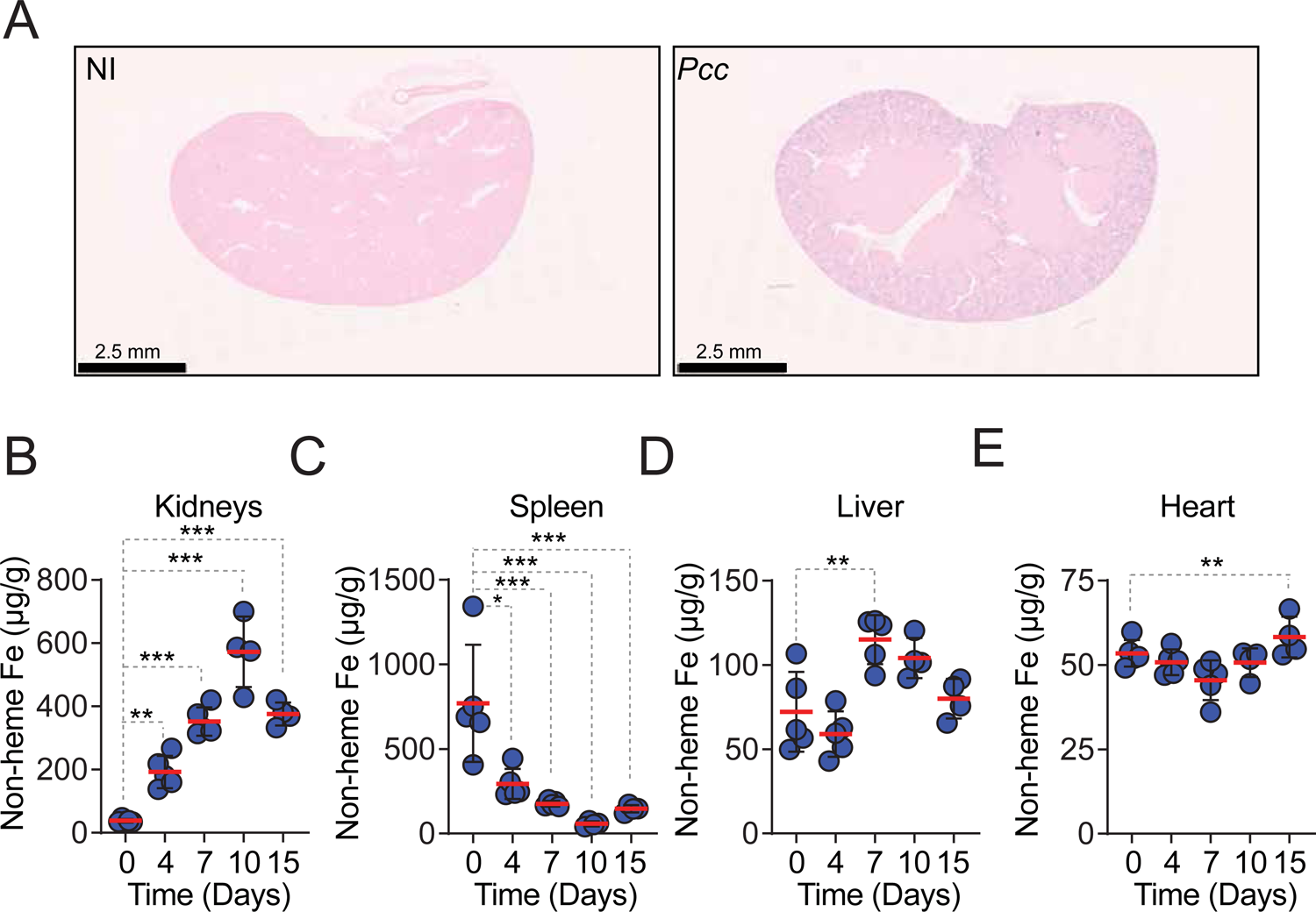
Peripheral Fe storage during *Pcc* infection. **A)** Staining of non-heme Fe using Perl’s Prussian Blue stain, in the kidneys of C57BL/6 mice non-infected (NI) or 7 days after *Pcc* infection (*Pcc*). Images are representative of 3 mice, from the same experiment as in (*Fig. 1B*). Scale bars: 2,5 mm. **B-E)** Non-heme iron *per* g of **B)** kidney, **C)** spleen, **D)** liver and **E)** heart of C57BL/6 mice non-infected (Day 0) or at different days after *Pcc* infection. Data represented as mean (red bar) ± SD, in one experiment. Circles in (B-E) represent individual mice. P values in (B-E) calculated by one-way ANOVA, NS: not significant; *: P<0.05; **: P<0.01; ***: P<0.001.

**Extended Data Figure 2.**
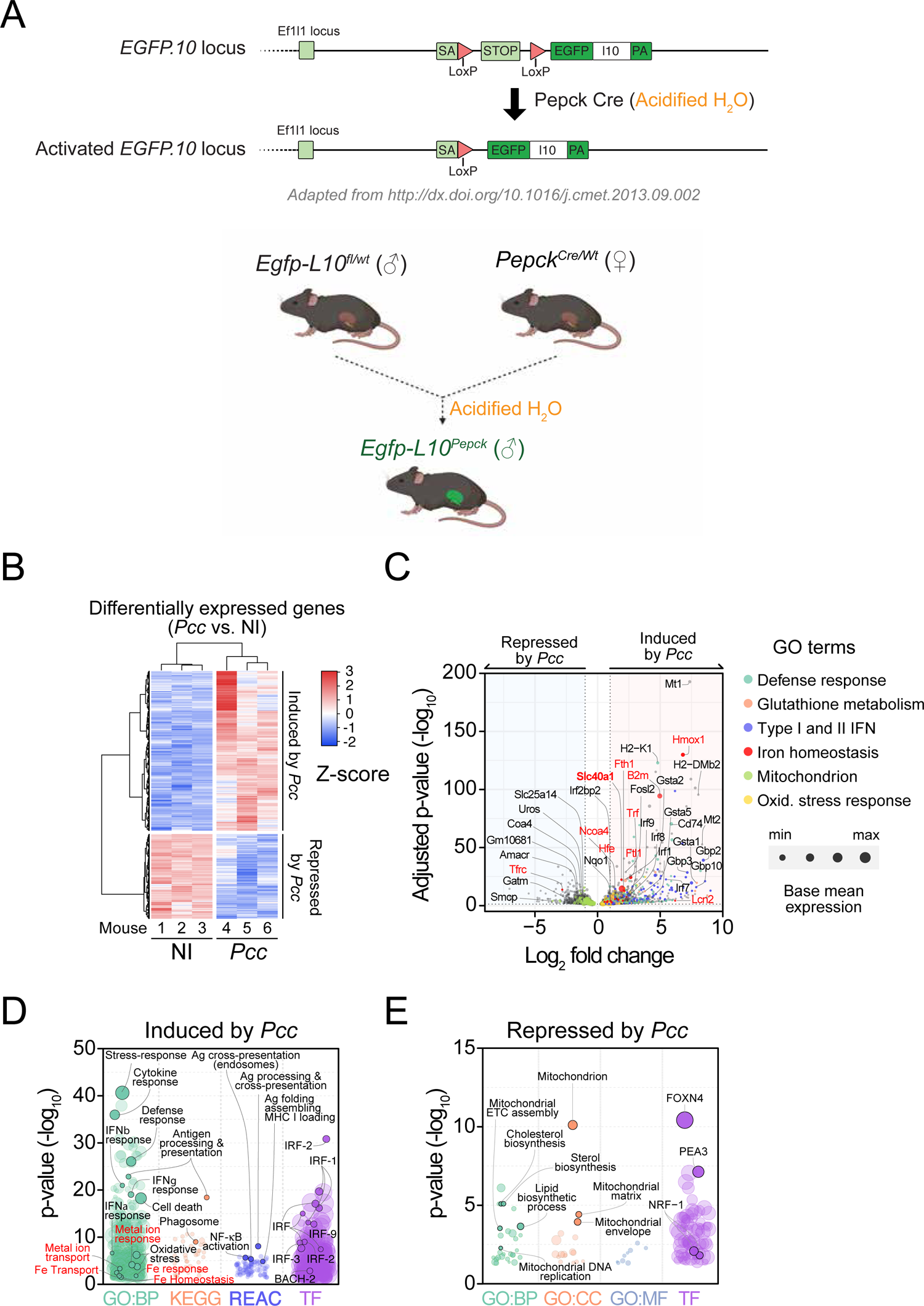
Translating Ribosome Affinity Purification (TRAP) from *Egfp-L10*^Pepck^ mice. **A)** Generation of *Egfp-L10*^Pepck^ mice expressing EGFP-tagged L10 ribosomal subunit specifically in RPTEC. Male *Egfp-L10^fl/wt^* mice, expressing a knocked in EGFP-tagged L10 ribosomal subunit, were crossed with female *Pepck^Cre/Wt^* mice, expressing a Cre recombinase under the control of a modified *Pepck* promoter driving Cre expression in RPTEC upon exposure to acidified drinking water. **B)** Heatmap and unsupervised clustering analysis of differentially expressed genes from RPTEC of *Egfp-L10*^Pepck^ mice, either non-infected (NI, N=3) mice or 7 days after *Pcc* infection (*Pcc*, N=3). Data from one experiment, displayed as scaled log transformed transcripts (Z-score). Same mice as in (*Fig. 1D*) (*see Tables S1, 2*). **C)** Volcano plot of differentially expressed genes, overlaid with color-labeled representative enriched gene ontologies for pairwise comparison between *Pcc*-infected and non-infected *Egfp-l10^Pepck^* mice **D,E)** Functional enrichment analysis of genes **D**) up-regulated and **E**) down-regulated in *Pcc*-infected *vs.* non-infected *Egfp-l10^Pepck^* mice. Representative ontologies color-coded. Databases: GO:BP=Gene ontology: Biological processes; GO:CC=Gene ontology: Cellular components; GO:MF=Gene ontology: Molecular functions; TF=Transfac, Transcription factor binding sites. KEGG: Kyoto Encyclopedia of Genes and Genomes, REAC: Reactome Pathway Database.

**Extended Data Figure 3.**
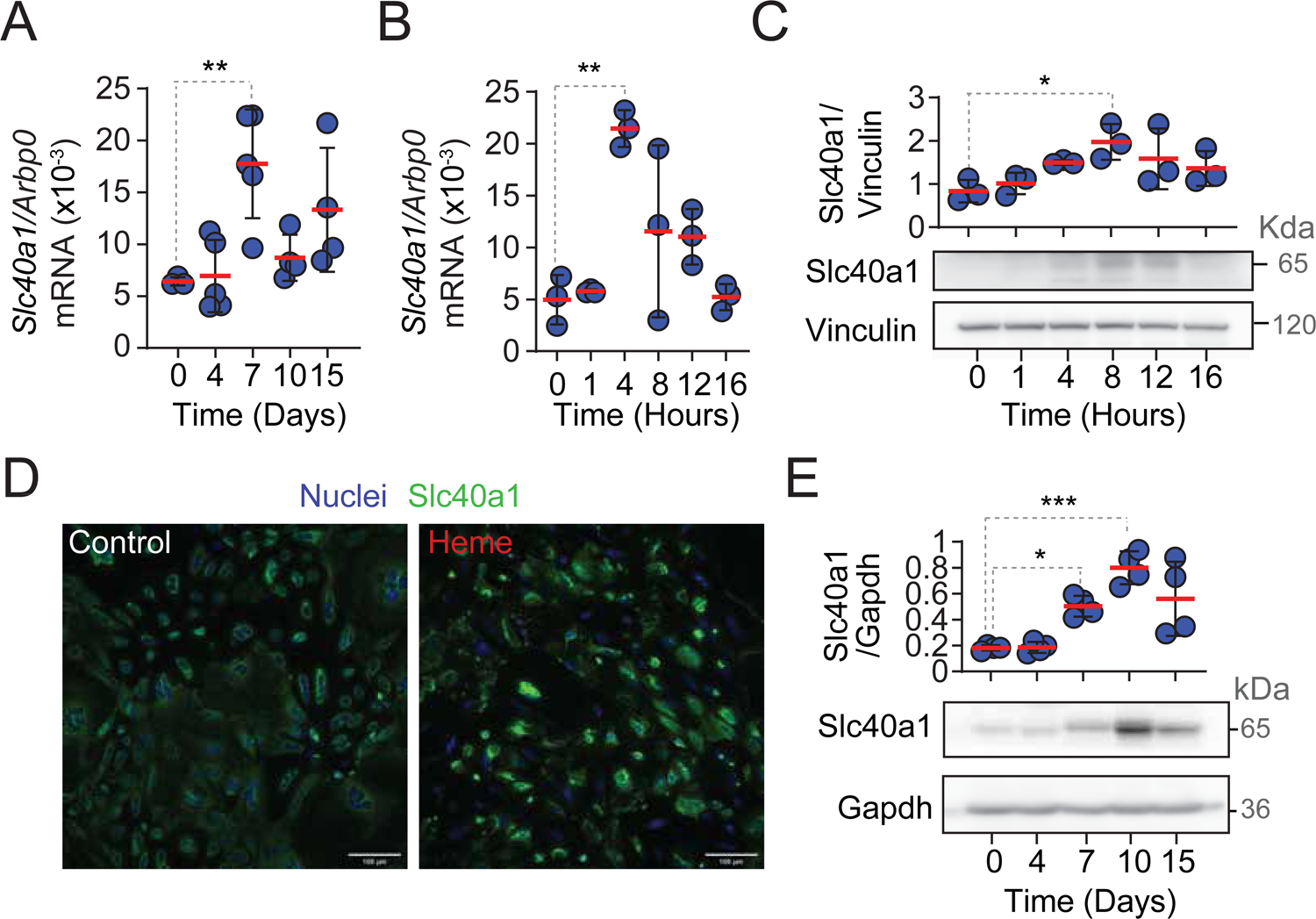
Induction of Slc40a1 in kidney and RPTEC. **A)** Relative expression of *Slc40a1* mRNA in the same experiment as in (*Fig. 1B*). Data represented as mean (red bar) ± SD. **B)** Induction of *Slc40a1* mRNA and **C)** Slc40a1 protein expression in RPTEC isolated from C57BL/6 mice exposed *in vitro* to heme (10µM). Data represented as mean ± SD, from two experiments. Blot in (C) is representative of 3 wells *per* time-point. The upper panel in (C) shows relative quantification of Slc40a1 expression, normalized to Gapdh. **C)** Expression of Slc40a1 in control or heme treated (10µM, 8h) RPTEC isolated from C57BL/6 mice. The immunofluorescence image is representative of 3 wells. Green: Slc40a1, blue: DAPI. Scale bars: 100µm. **D)** Slc40a1 protein expression in kidney lysates from *Pcc*-infected C57BL/6 mice before (Day 0) or after *Pcc* infection. The blot is representative of 4 animals *per* time-point. Upper panel shows relative densitometry quantification of Slc40a1/Gapdh represented as mean ± SD from one out two experiments with similar trend. Circles correspond to individual mice. Circles in (A, E) represent individual mice and in (B,C) individual wells. P values in (A,B,C,E) calculated by one-way ANOVA, NS: not significant; *: P<0.05; **: P<0.01; ***: P<0.001.

**Extended Data Figure 4.**
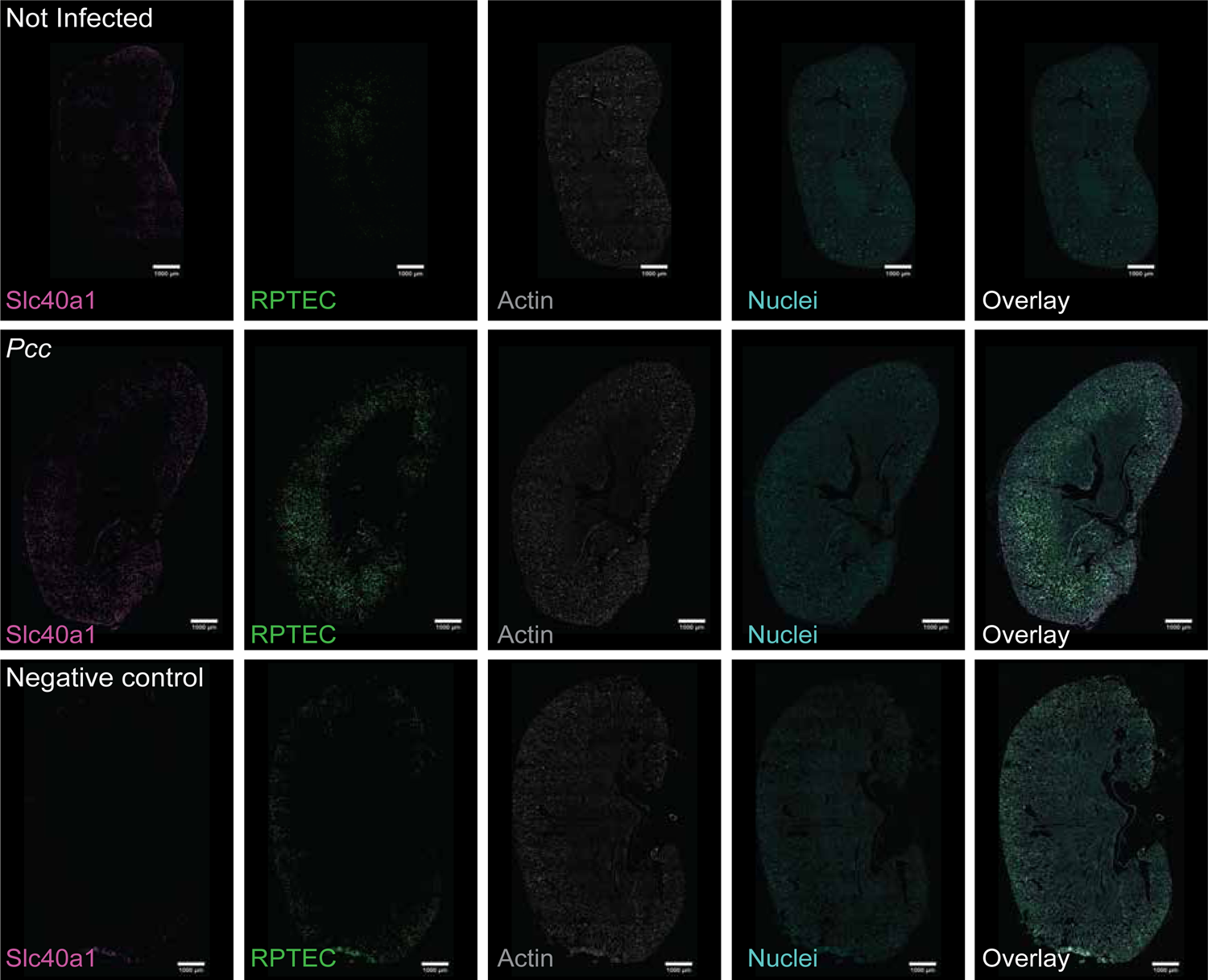
Renal Slc40A1 expression during *Pcc* infection. Immunofluorescence imaging of kidney sections from control (NI, non-infected) *vs. Pcc*-infected (day 7) *Egfp-L10^Pepck^* mice, expressing GFP specifically in RPTEC (green). Slc40a1 (magenta) was stained using a polyclonal antibody, DNA (nuclei) with DAPI (cyan) and actin (grey) was labeled using fluorescent-conjugated Phalloidin. Same samples as in (*Fig. 1F*). Negative control was performed using kidney sections from *Slc40a1*^fl/fl^ mice, omitting the primary antibody. Images are representative of 3 animals *per* group from 2 independent experiments. Scale bars: 1mm.

**Extended Data Figure 5.**
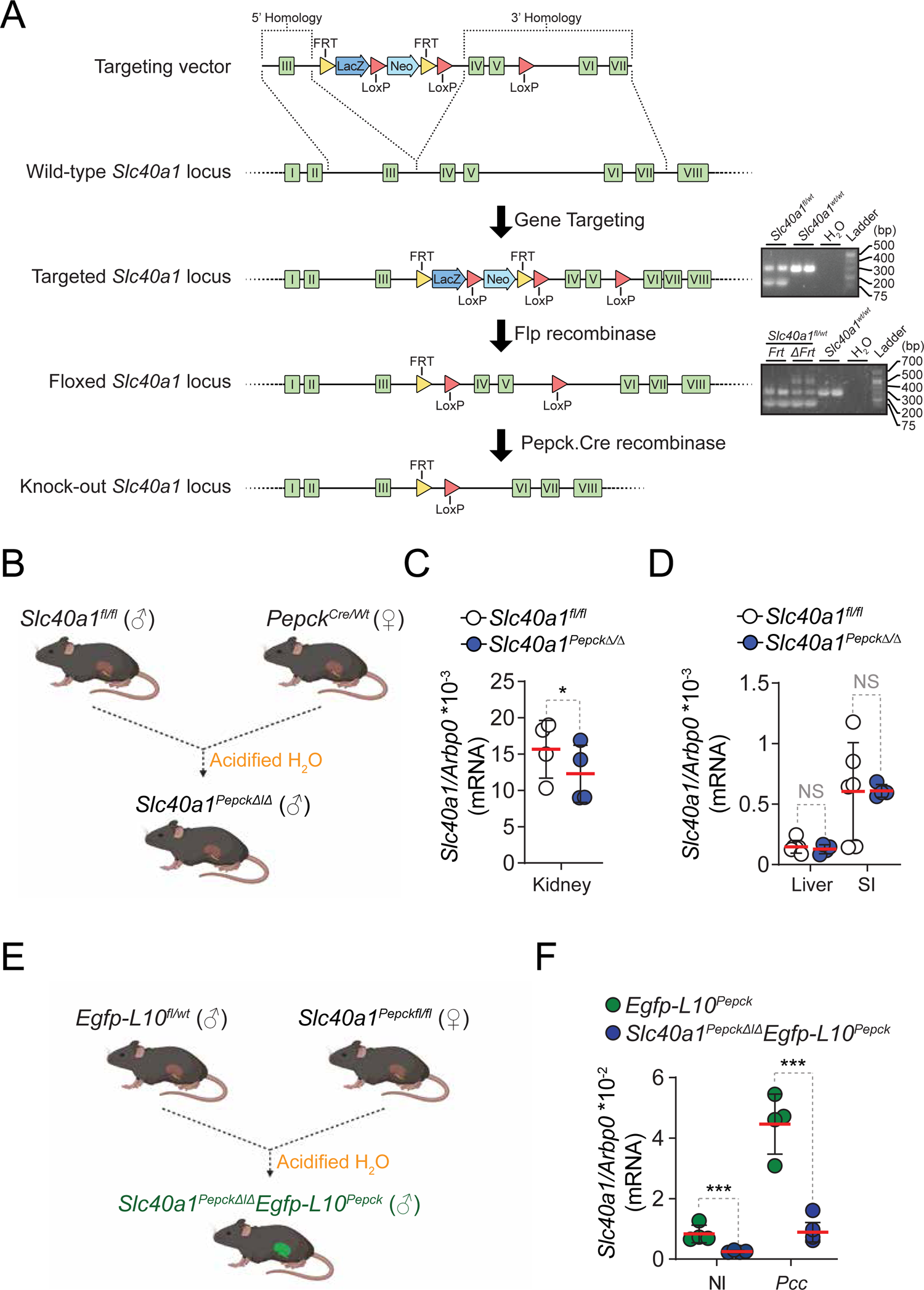
*Slc40a1^PepckΔ/Δ^Egfp-L10^Pepck^* mice. **A, B**) Sperm containing the full Knockout-First mutant of the *Slc40a1* allele floxed, with flox sequences flanking exons 4 and 5, was used to fertilize oocytes from C57BL/6 mice. Off-spring was crossed with C57BL/6 Flp recombinase mice to delete the *frt*-flanked sequence and generate *Slc40a1^fl/fl^* mice, which were crossed with *Pepck^Cre^* mice to generate *Slc40a1^Pepckfl/fl^* mice, where the floxed *Slc40a1^fl/fl^* allele was specifically deleted in RPTEC from *Slc40a1^Pepck^*^Δ/Δ^ mice after exposure to acidified drinking water. The breeding strategy consisted of crossing male *Slc40a1^fl/fl^* with female *Slc40a1^Pepckfl/fl^* mice to generate male *Slc40a1^Pepckfl/fl^* and control littermate male *Slc40a1^fl/fl^* mice to use in experiments after exposure to acidified drinking water. **C, D**) Quantification of *Slc40a1* mRNA in **C**) kidneys, **D**) liver and small intestine (SI) of *Slc40a1^fl/fl^ vs. Slc40a1^PepckΔ/Δ^* mice, by qRT-PCR. Data represented as mean (red bar) ± SD. Circles represent individual mice (N=4-6 *per* genotype). **E)** *Slc40a1^PepckΔ/Δ^Egfp-L10^Pepck^* mice expressing an EGFP-tagged L10 ribosomal subunit specifically in RPTEC were generated by crossing the *Slc40a1^Pepckfl/fl^* mice produced in (A) with *Egfp-L10^fl/wt^Slc40a1^fl/fl^* mice (*See Fig. S2*). **F)** *Slc40a1* mRNA expression in kidneys from *Egfp-L10^Pepck^* and *Slc40a1^PepckΔ/Δ^Egfp-L10^Pepck^* mice not infected (NI) or 7 days after *Pcc* infection. Data represented as mean (red bar) ± SD, from 4 independent experiments. Circles represent individual mice (N= 4-8 *per* group). P values in (C) determined using Paired t test and in (D, F) using one-way ANOVA, NS: not significant; *: P<0.05; **: P<0.01; ***: P<0.001.

**Extended Data Figure 6.**
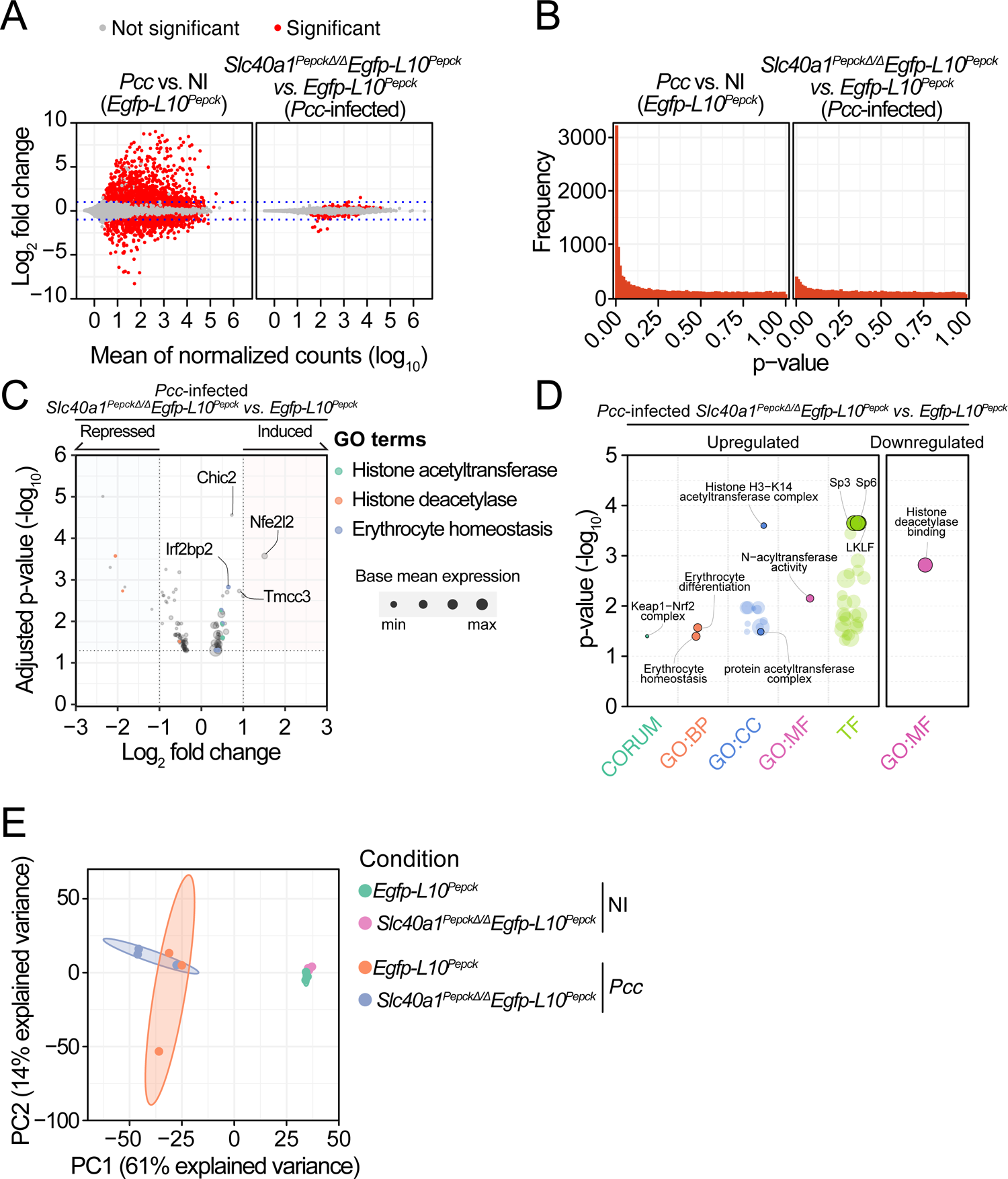
Effect of *Slc40a1* deletion in RPTEC. **A-E)** Active transcriptome analysis of RPTEC from *Pcc*-infected and non-infected *Egfp-L10^Pepck^* mice (expressing *Slc40a1* in RPTEC) or *Slc40a1^PepckΔ/Δ^Egfp-L10^Pepck^* mice (*Slc40a1* deleted in RPTEC). **A)** MA (log ratio-mean average) plots of gene expression differences between *Pcc*-infected *vs.* non-infected (NI) *Egfp-L10^Pepck^* mice (left panel) or *Pcc*-infected *Slc40a1^PepckΔ/Δ^Egfp-L10^Pepck^* vs. *Pcc*-infected *Egfp-L10^Pepck^* mice (right panel). **B)** Adjusted p-value frequency histogram plots calculated for each gene, categorizing gene expression differences between RPTEC from *Pcc*-infected *vs*. non-infected (NI) *Egfp-L10^Pepck^* mice (left panel) or *Pcc*-infected *Slc40a1^PepckΔ/Δ^Egfp-L10^Pepck^* vs. *Pcc*-infected *Egfp-L10^Pepck^* mice (right panel). **C)** Volcano plot of differentially expressed genes, overlaid with representative enriched gene ontologies for pairwise comparison between *Pcc*-infected *Slc40a1^PepckΔ/Δ^Egfp-L10^Pepck^* vs. *Egfp-L10^Pepck^* mice. Representative differentially expressed genes labeled. **D)** Functional enrichment analysis of genes up-(left panel) and downregulated (right panel) in RPTEC from *Pcc*-infected *Slc40a1^PepckΔ/Δ^Egfp-L10^Pepck^* vs. *Pcc*-infected *Egfp-L10^Pepck^* mice. Representative ontologies highlighted. Databases: CORUM = comprehensive resource of mammalian protein complexes; GO:BP=Gene ontology: Biological processes; GO:CC=Gene ontology: cellular component; GO:MF=Gene ontology: molecular function; TF=Transfac, Transcription factor binding sites. **E)** Principal component analysis (PCA) of mRNAseq data from non-infected (NI) and *Pcc*-infected, *Slc40a1^PepckΔ/Δ^Egfp-L10^Pepck^* and *Egfp-L10^Pepck^* mice (N=3 per condition), and 90% confidence ellipses for each condition. (*see Tables S1, 3*).

**Extended Data Figure 7.**
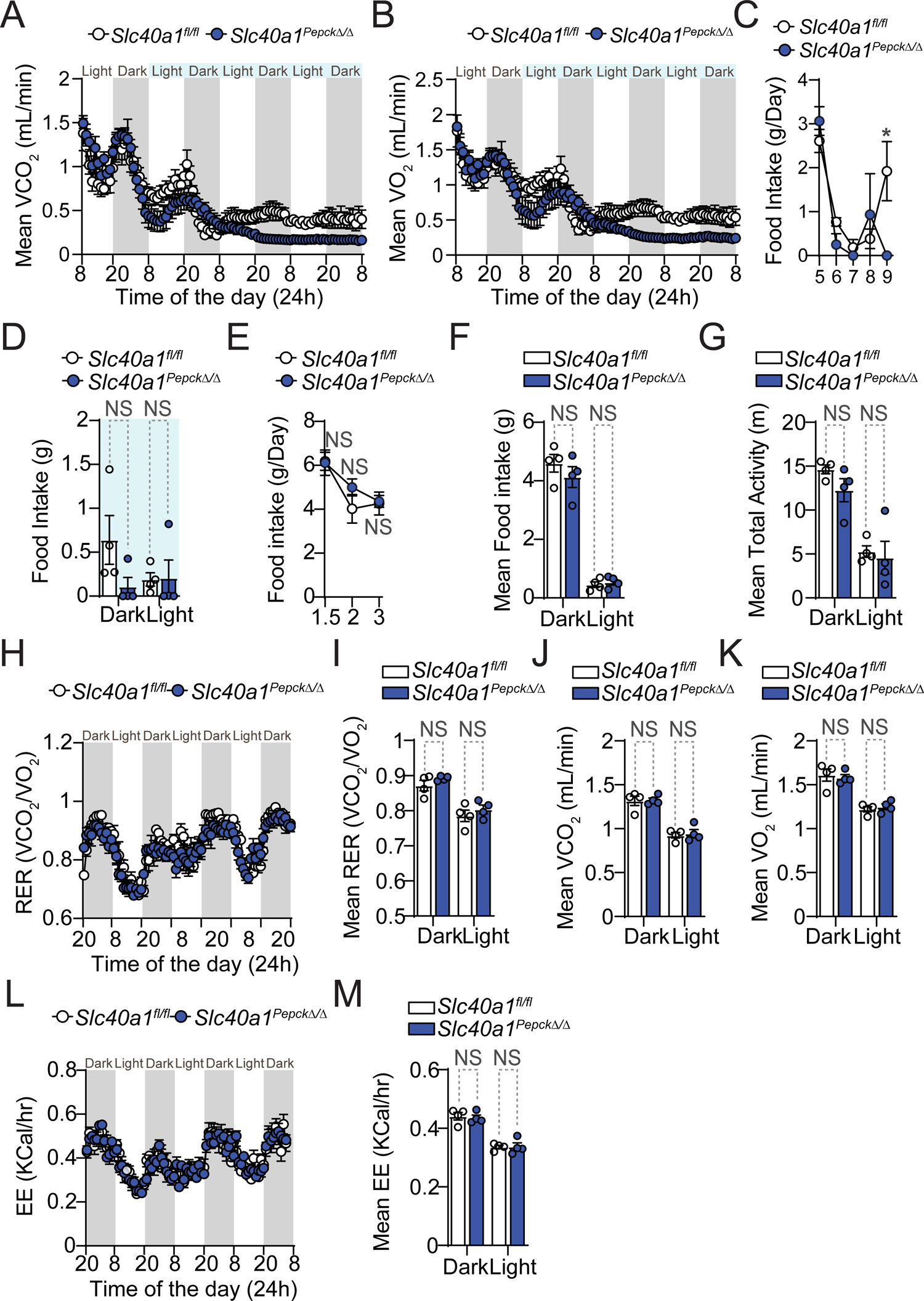
RPTEC *Slc40a1* supports organismal metabolic reprogramming. **A-D)** Synchronized metabolic and behavioral quantification from 5 to 10 days after *Pcc* infection in *Slc40a1^Pepck^*^Δ/Δ^ (N=4, blue circles) and *Slc40a1*^fl/fl^ (N=4, white circles) mice (same experiment as *Fig. 2D-K*). Daily **A)** VCO_2_, **B)** VO_2_ and **C)** Food intake. **D)** mean food intake ± SEM in the period indicated (light blue in A,B), segregated into daily light/dark cycle. **E-M)** Steady-state synchronized metabolic and behavioral quantification from *Slc40a1^Pepck^*^Δ/Δ^ (N=4, blue circles) and *Slc40a1*^fl/fl^ (N=4, white circles) mice in one experiment. **E)** Daily food intake. **F)** Mean ± SEM of daily averages of food intake. **G)** Mean ± SEM of daily averages of total activity. **H)** Daily respiratory exchange ratio (RER). **I-K)** Mean ± SEM of daily averages of **I)** RER, **J)** VCO_2_ and **K)** VO_2_, corresponding to the period represented in (H). Data is segregated into daily light/dark cycles. **L)** Daily energy expenditure (EE). **M)** Mean ± SEM of daily averages of EE in the period represented in (H), segregated into daily light/dark cycle. In (F,G,I-K,M) circles represent average of 3 days of recording of individual mice. P values in (C-G, I-K and M) determined using two-way ANOVA. NS: not significant.

**Extended Data Figure 8.**
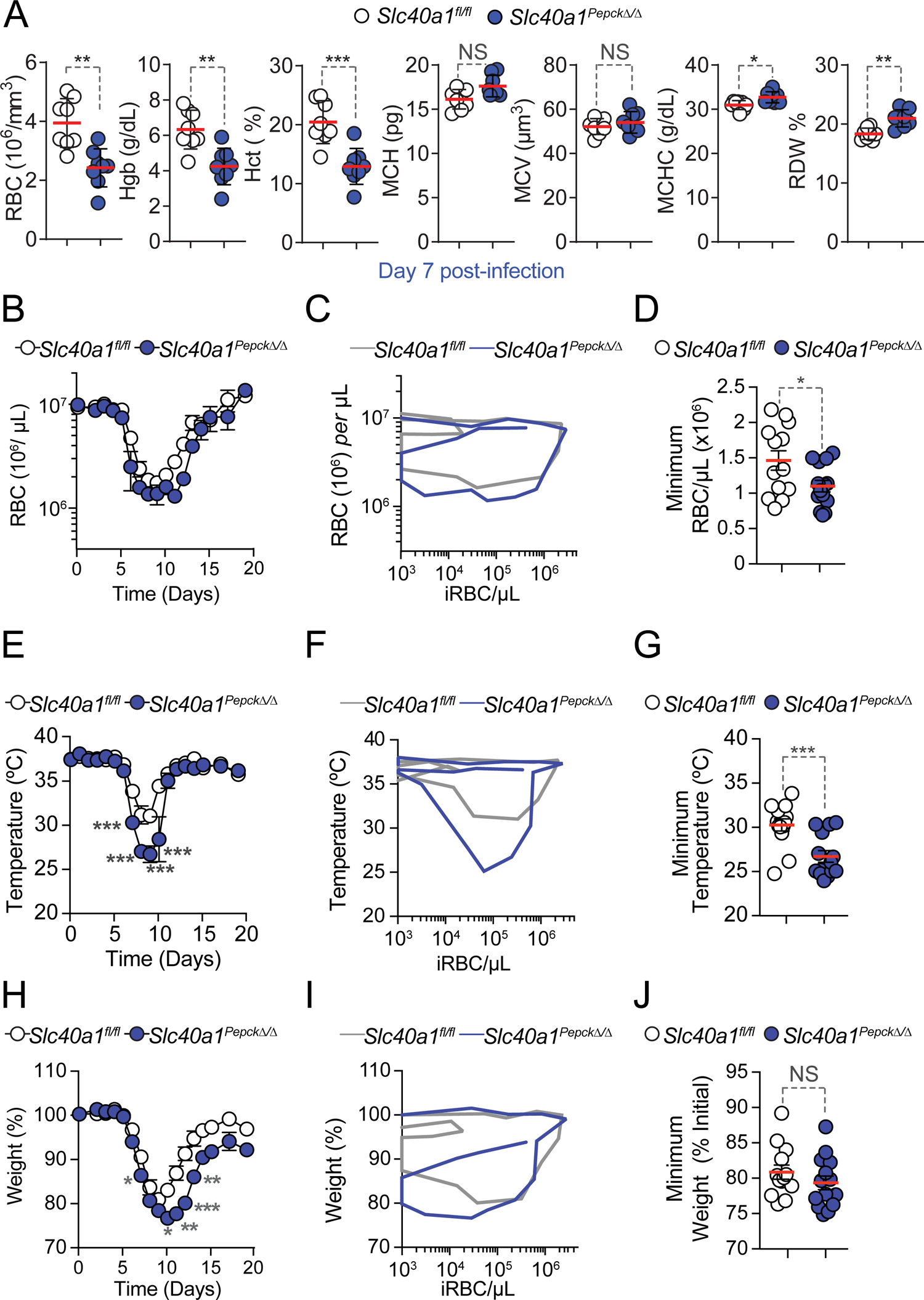
RPTEC *Slc40a1* supports RBC homeostasis. **A)** Hemograms of *Slc40a1^Pepck^*^Δ/Δ^ (N=8) and *Slc40a1*^fl/fl^ (N=8) mice, 7 days after *Pcc* infection. Data represented as mean (red bar) ± SD, derived from two independent experiments with similar trend. RBC: red blood cell count; Hb: hemoglobin concentration; Hct: hematocrit; MCH: mean corpuscular hemoglobin; MCH: mean corpuscular volume; MCHC: mean corpuscular hemoglobin concentration; RDW: RBC distribution width. **B-J)** Disease parameters in *Pcc* infected *Slc40a1^Pepck^*^Δ/Δ^ (N=14) and *Slc40a1*^fl/fl^ (N=13) mice. Data from the same experiments as in (*Fig. 2A-C*). **B)** Mean circulating RBC numbers ± SEM. **C)** Disease trajectories established by the relationship of the median number of circulating RBC *per* µL *vs.* median number of infected RBC (iRBC) per µL (i.e, pathogen load). **D)** Minimum number of circulating RBC *per* µL (mean ± SEM) during the course of *Pcc* infection. **E)** Mean temperature ± SEM. **F)** Disease trajectories established by the relationship of median body temperature *vs.* the median number of infected RBC (iRBC) *per* µL (*i.e.,* pathogen load). **G)** Minimum temperature (mean ± SEM) during the course of *Pcc* infection. **H)** Mean percentage of initial body weight ± SEM. **I)** Disease trajectories established by the relationship between the median percentage of initial body weight *vs.* the median number of infected RBC (iRBC) per µL (*i.e.,* pathogen load). **J)** Minimum percentage of initial body weight (mean ± SEM) during the course of *Pcc* infection. Circles in (A, D, G and J) correspond to individual mice. P values in (A, D, G and J) determined using Mann Whitney test and in (B, E, and H) using one-way ANOVA, NS: not significant; *: P<0.05; **: P<0.01; ***: P<0.001.

**Extended Data Figure 9.**
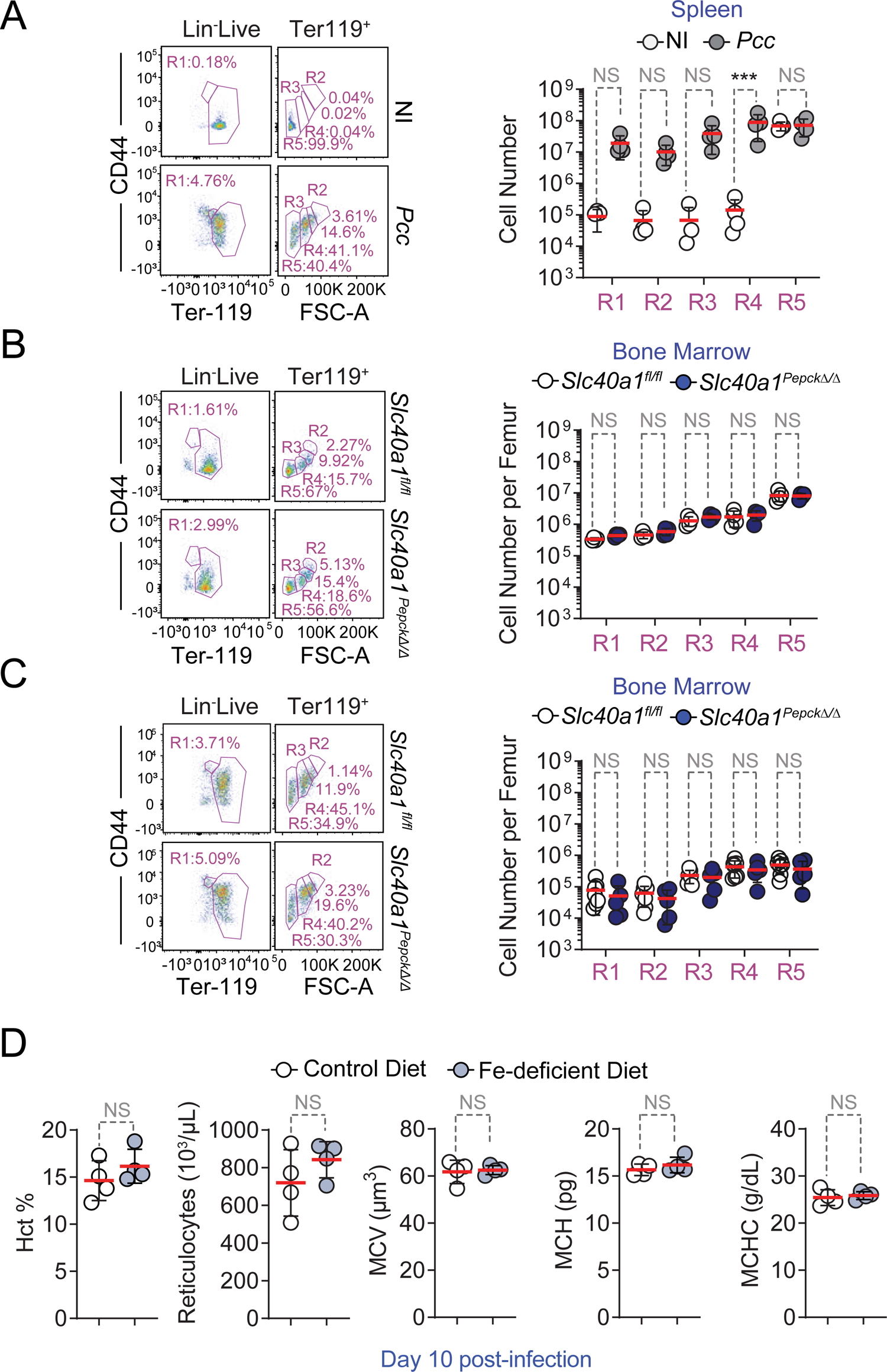
RPTEC *Slc40a1* promotes erythropoiesis. **A)** Extramedullary erythropoiesis in the spleen of control C57BL/6 mice (*i.e.,* non-infected; NI; N=4) and 10 days after *Pcc* infection (N=4). Data from one experiment. Left panel is a representative FACS plot and gating strategy based on size, CD44 and Ter119 expression used to identify different RBC developmental stages. Full gating strategy in *Figure S_1.* Right panel represents the mean (red bar) ± SD number of each cell population. **B-C)** Medullary erythropoiesis in **B)** non-infected *Slc40a1^Pepck^*^Δ/Δ^ and *Slc40a1*^fl/fl^ mice (N=4 *per* genotype) and **C)** 10 days after *Pcc* infection in *Slc40a1^Pepck^*^Δ/Δ^ (N=5) and *Slc40a1*^fl/fl^ (N=7) mice. Data from the same mice as in (*Fig. 2L*). Left panels are representative FACS plots and gating strategies, based on size, CD44 and Ter119 expression, used to identify different RBC developmental stages. Full gating strategy in *Fig. S_1.* Right panels are the means (red bars) ± SD of the number of each cell population. **D)** Hemogram analysis of *Pcc*-infected C57BL/6 mice, fed on Fe-deficient (N=4) vs. control chow (N=4) diet, 10 days post infection. Data represented as mean (red bar) ± SD, derived from 1 experiment (same mice as in *Fig. 3G-I*). Hct: hematocrit; MCH: mean corpuscular hemoglobin; MCH: mean corpuscular volume; MCHC: mean corpuscular hemoglobin concentration; RDW: RBC distribution width. Circles in (A-D) correspond to individual mice. P values in (A-C) determined using one-way ANOVA and in (D) using Mann Whitney test. NS: not significant; ***: P<0.001.

**Extended Data Figure 10.**
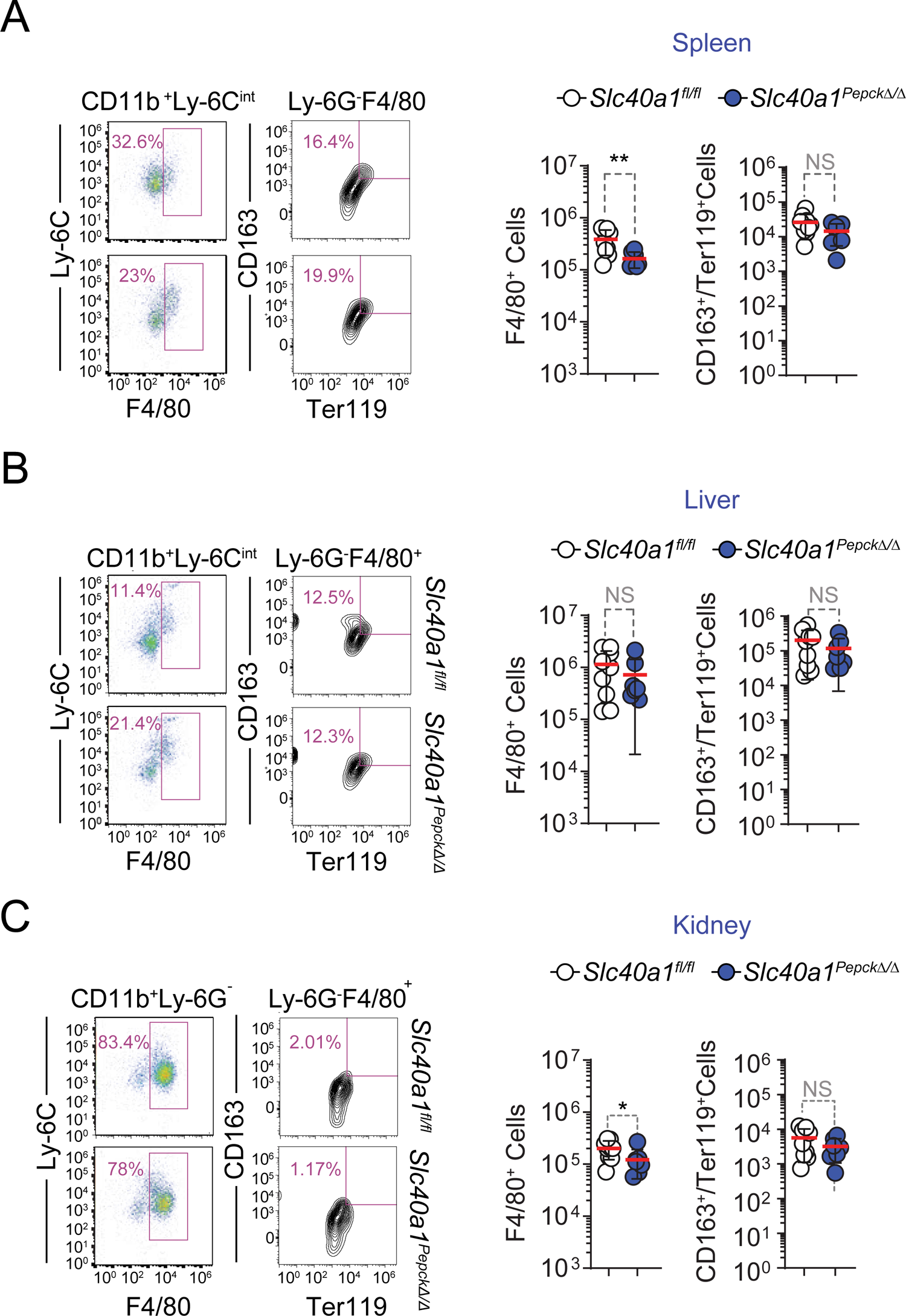
RPTEC Slc40a1 does not promote erythrophagocytosis. Erythrophagocytosis in **A)** spleen, **B)** liver and **C)** kidneys of *Slc40a1^Pepck^*^Δ/Δ^ (N=7) and *Slc40a1*^fl/fl^ (N=9) mice, 7 days after *Pcc* infection. Data from two independent experiments with similar trend. Left panels are representative FACS plots of the gating strategy used to identify erythrophagocytic (CD11b^+^, Ly-6C^int^, Ly-6G^-^, F4/80^+^, CD163^+^) macrophages engulfing RBC (intracellular Ter119^+^). Full gating strategy in *Figure S_3.* Right panels are the quantification of cell numbers represented as mean (red bar) ± SD, whereby circles represent individual mice. P values in (A-C) were determined using Mann Whitney test, NS: not significant; *: P<0.05; **: P<0.01.

**Extended Data Figure 11.**
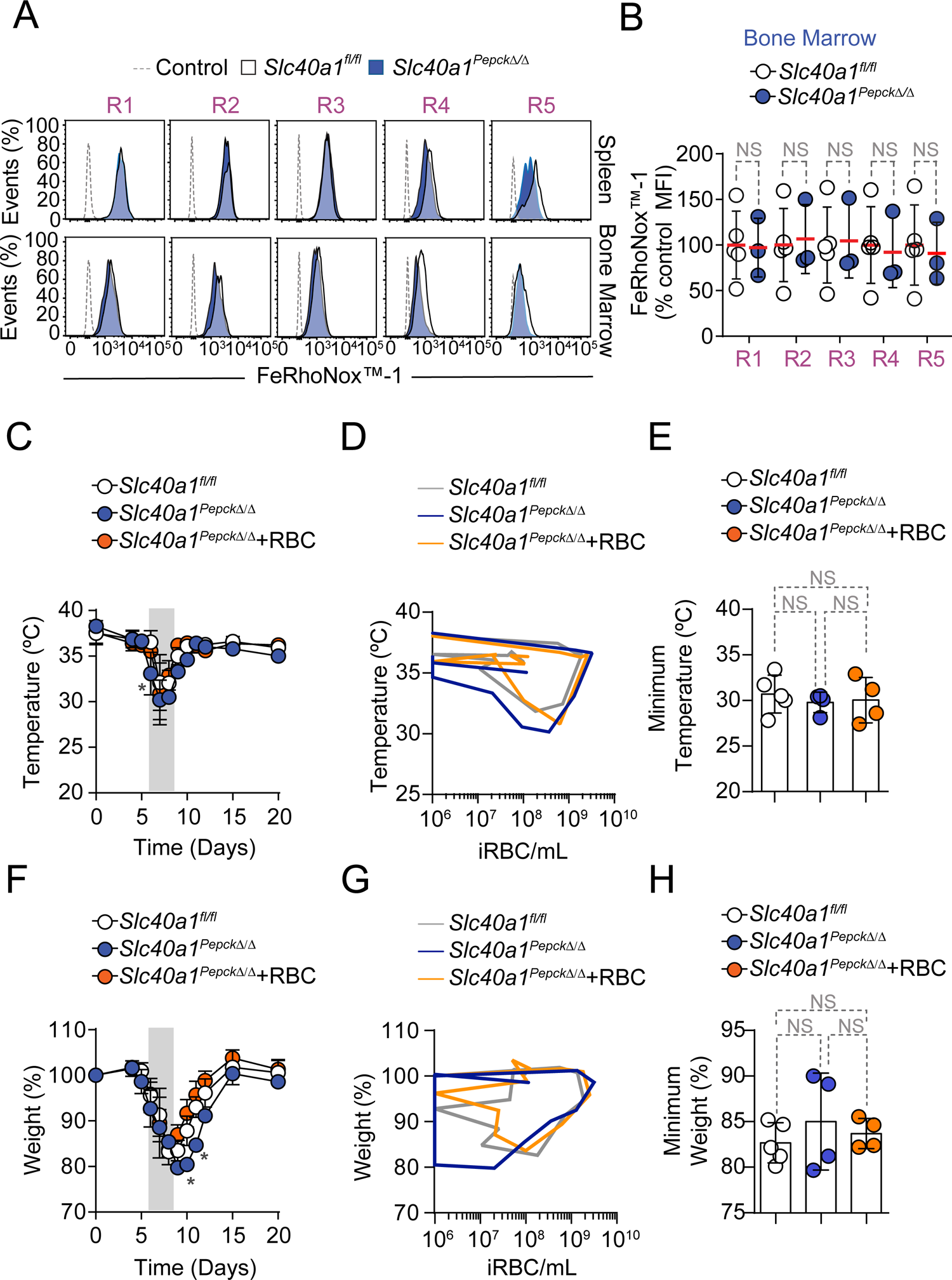
Protective effect of RBC transfusion. **A)** Representative flow cytometry histogram of Fe^2+^ (FeRhoNox^TM^-1) staining in splenic (top panels) and bone marrow (bottom panels) erythroblasts from *Pcc*-infected *Slc40a1^Pepck^*^Δ/Δ^ (N=3) and *Slc40a1^fl/fl^* (N=5) mice from two independent experiments with similar trend. Cell populations were identified based on the gating strategy detailed in (*Fig. S_3*). Dashed line (Control) corresponds to basal signal in C57BL/6J wild type cells without the FeRhoNox^TM^-1 probe. **B)** Relative quantification of Fe^2+^ (FeRhoNox^TM^-1) staining from bone marrow, normalized to the average mean fluorescence intensity (MFI) of each erythroblast stage (*i.e.,* R1-5) in *Pcc*-infected *Slc40a1^fl/fl^* mice, represented as mean (red bar) ± SD. (**C-H**) *Pcc*-infected *Slc40a1^Pepck^*^Δ/Δ^ mice received naïve fresh RBC (8×10^8^, i.p., N=4) or vehicle (PBS) as control (N=4). *Pcc*-infected *Slc40a1*^fl/fl^ receiving vehicle (PBS) were used as an additional control (N=4). Data pooled from two out of 3 independent experiments, with similar trend (same mice as in *Fig. 3-K,L*). **C)** Mean temperature ± SD. **D)** Disease trajectories established by the relationship of median body temperature *vs.* median number of infected RBC (iRBC) *per* µL (*i.e.,* pathogen load). **E)** Minimum temperature (mean ± SD) during the course of *Pcc* infection. **F)** Mean percentage of initial body weight ± SD. **G)** Disease trajectories established by the relationship between the median percentage of initial body weight *vs.* the median number of infected RBC (iRBC) *per* µL (*i.e.,* pathogen load). **H** Minimum percentage of initial body weight (mean ± SD) during the course of *Pcc* infection. Circles in (B,E,H) correspond to individual mice. P values in (B,E,H) were determined using one-way ANOVA and in (C,F) using two-way ANOVA. NS: not significant; *: P<0.05.

**Extended Data Figure 12.**
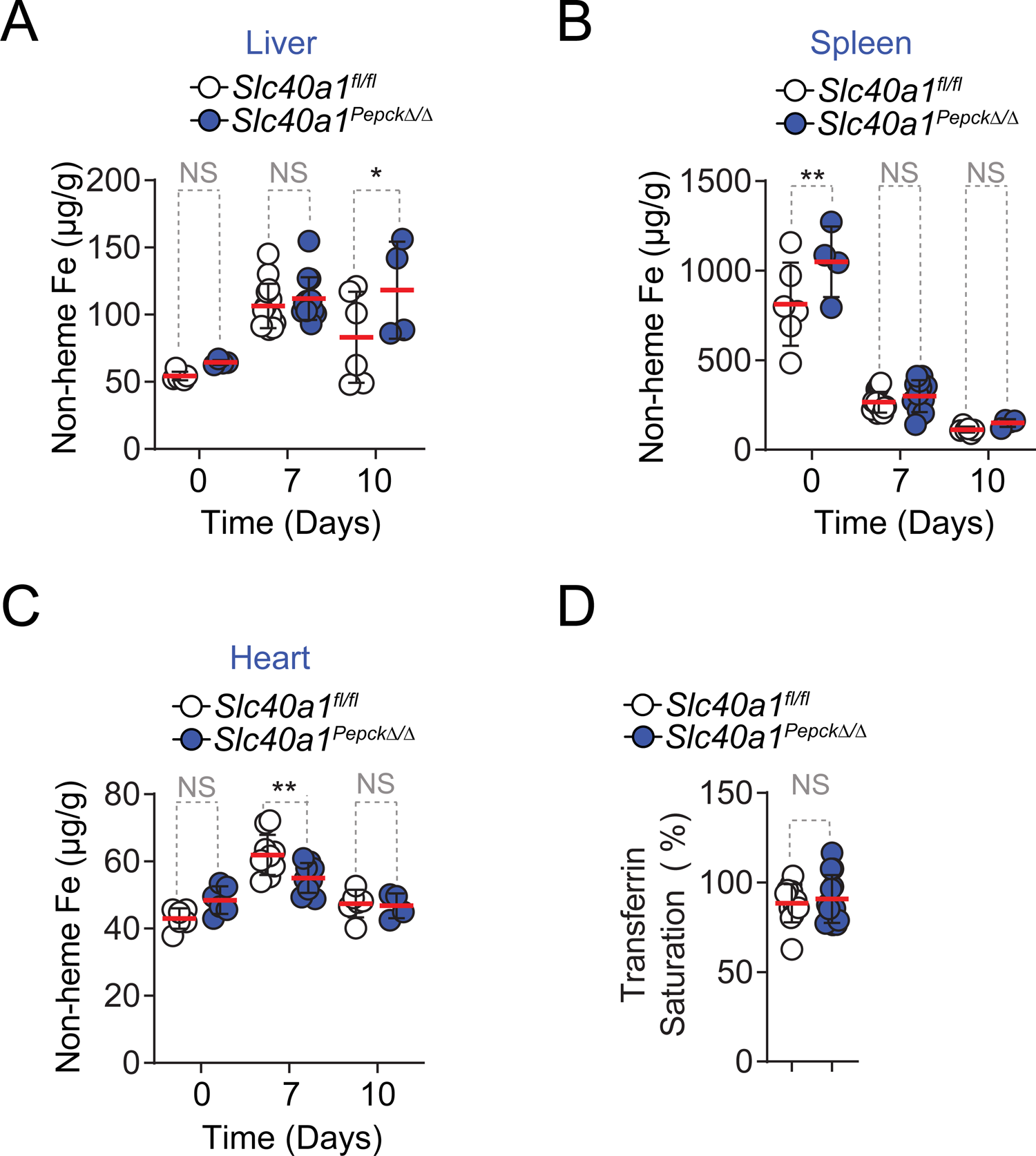
Tissue iron distribution in *Slc40a1^Pepck^*^Δ/Δ^ during *Pcc* infection. Non-heme Fe content in **A)** Liver, **B)** spleen, and **C)** heart of *Slc40a1^Pepck^*^Δ/Δ^ and *Slc40a1*^fl/fl^ mice, not infected (D0), at the peak (Day 7) or at recovery phase (Day 10) of *Pcc* infection. Data is represented as mean of Fe µg *per* g of tissue (red bar) ± SD (N=4-14 mice per time-point). Data from the same experiments as in (*Fig. 4A*). **D)** Transferrin saturation in the plasma of *Pcc*-infected *Slc40a1^Pepck^*^Δ/Δ^ (N=13) and *Slc40a1*^fl/fl^ (N=12) mice. Data is represented as mean (red bar) ± SD from four independent experiments. Circles in correspond to individual mice. P values in (A-C) were determined using one-way ANOVA and in (D) using Mann Whitney test. NS: not significant; *: P<0.05; **: P<0.01.

**Extended Data Figure 13.**
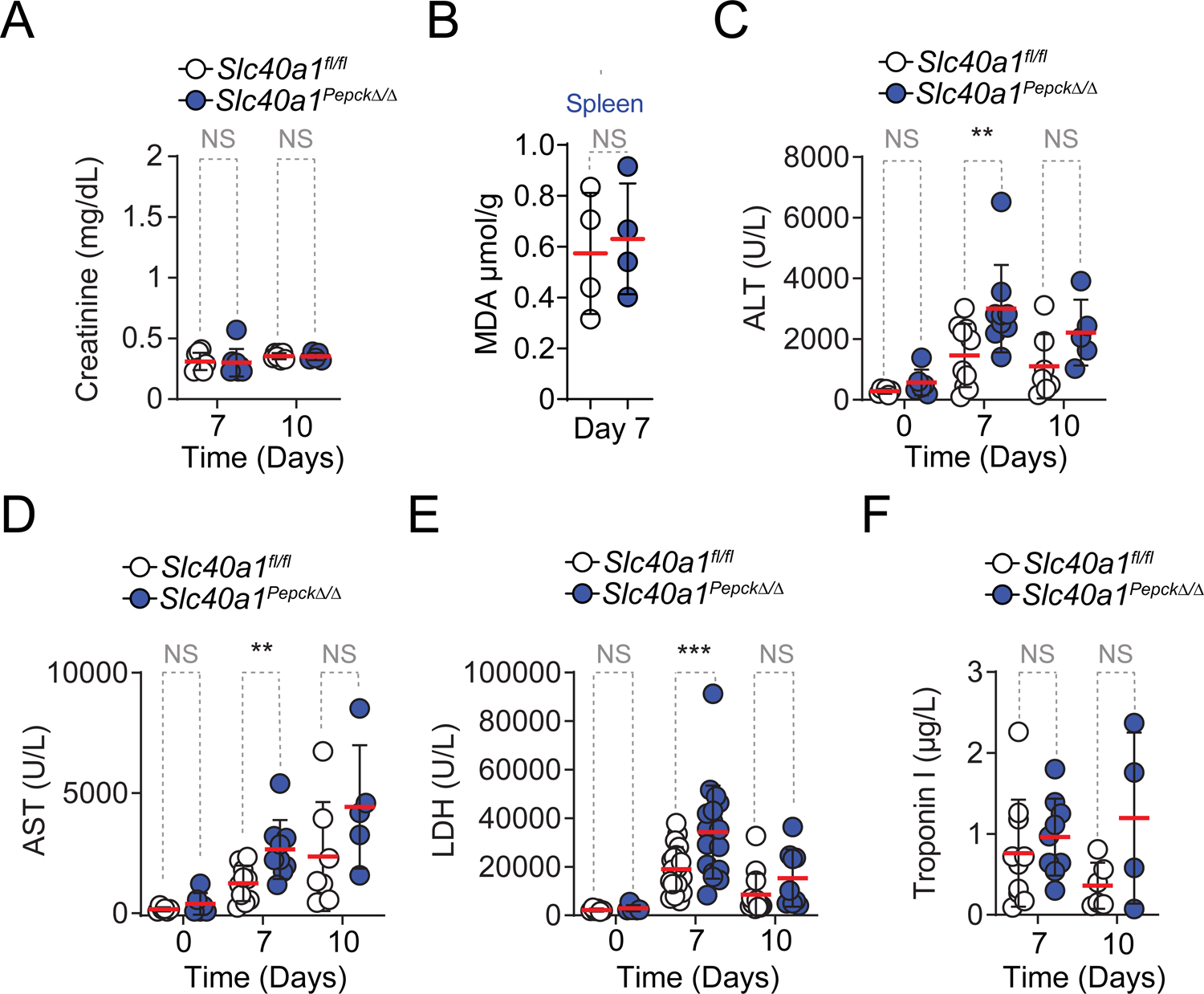
Impact of RPTEC Slc40a1 on serological markers of organ damage. **A)** Creatinine concentration in the plasma from *Slc40a1^Pepck^*^Δ/Δ^ and *Slc40a1*^fl/fl^ mice at the peak (Day 7) or recovery phase (Day 10) of *Pcc* infection, represented as mean (red bar) ± SD. (N=5-8 mice *per* time-point). Data from two independent experiments shown in (*Fig. 4A*). **B)** Quantification of the lipid peroxidation product malondialdehyde (MDA) in the spleen of *Slc40a1^Pepck^*^Δ/Δ^ and *Slc40a1*^fl/fl^ mice, 7 days after *Pcc* infection. Data represented as mean (red bar) ± SD, (N=4 mice *per* genotype), from one experiment. **C)** Alanine transaminase (ALT), **D)** Aspartate transaminase (AST), **E)** Lactate dehydrogenase and **F)** Troponin I concentrations in plasma from *Slc40a1^Pepck^*^Δ/Δ^ and *Slc40a1*^fl/fl^ mice, before infection (D0), at the peak (Day 7) or recovery phase (Day 10) of *Pcc* infection. Data is represented as mean (red bar) ± SD, (N=4-20 *mice per* time-point) from the same experiment as in (*Fig. 4E*). Circles represent individual mice. P values in (A, C-F) were determined using one-way ANOVA and in (B) using Mann Whitney test, NS: not significant; **: P<0.01, ***: P<0.001.

**Extended Data Figure 14.**
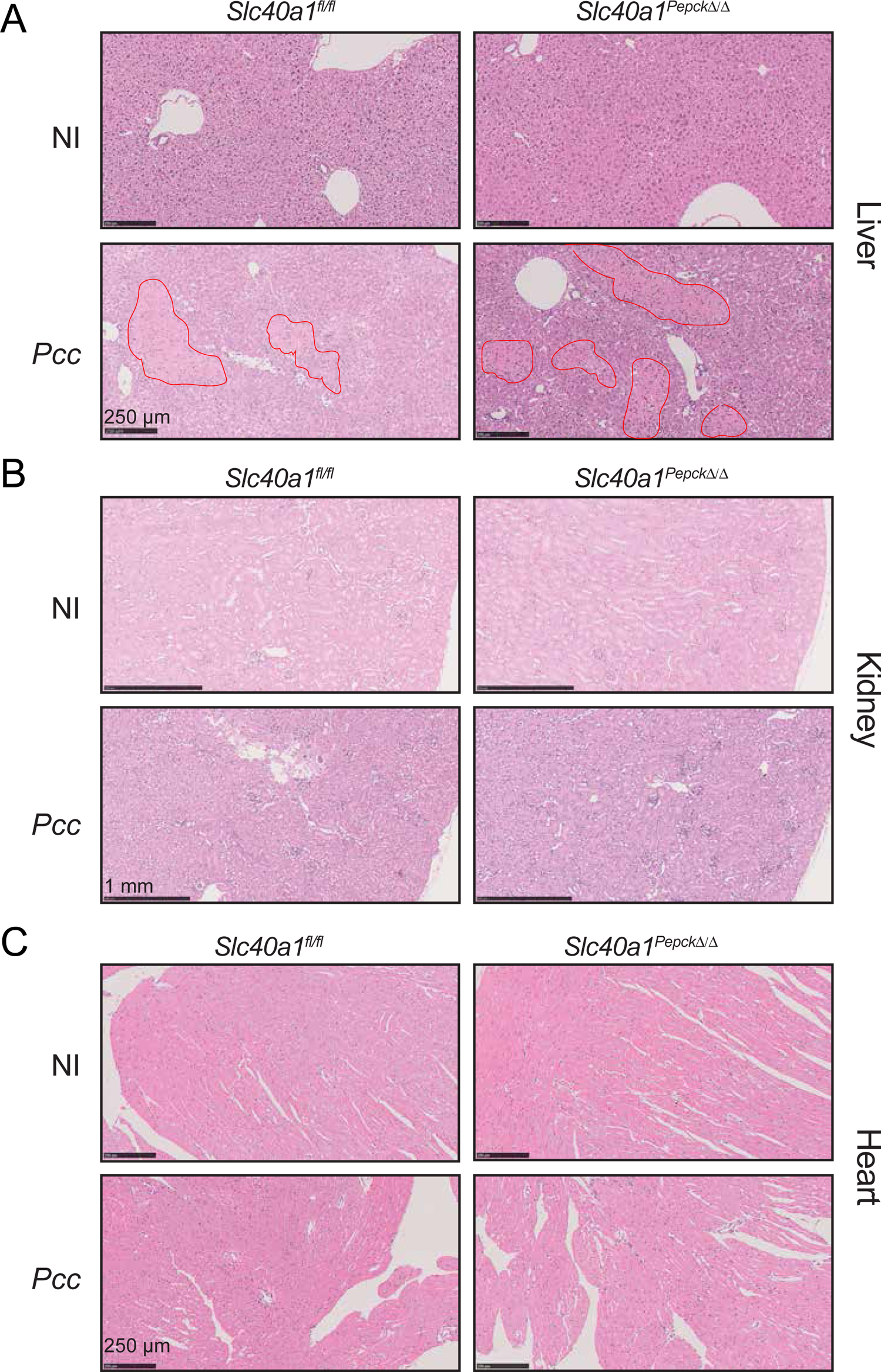
Impact of RPTEC Slc40a1 on organ damage. **A-C)** H&E-stained **A)** liver, **B)** kidney and **C)** heart from non-infected (NI; Day 0) and at the peak of *Pcc* infection (Day 7) in *Slc40a1^Pepck^*^Δ/Δ^ and *Slc40a1*^fl/fl^ mice. Images are representative of 3-5 animals in the same experiment as in (*Fig. 1B*). Red lines in (A) delineate liver necrotic areas.

**Extended Data Figure 15.**
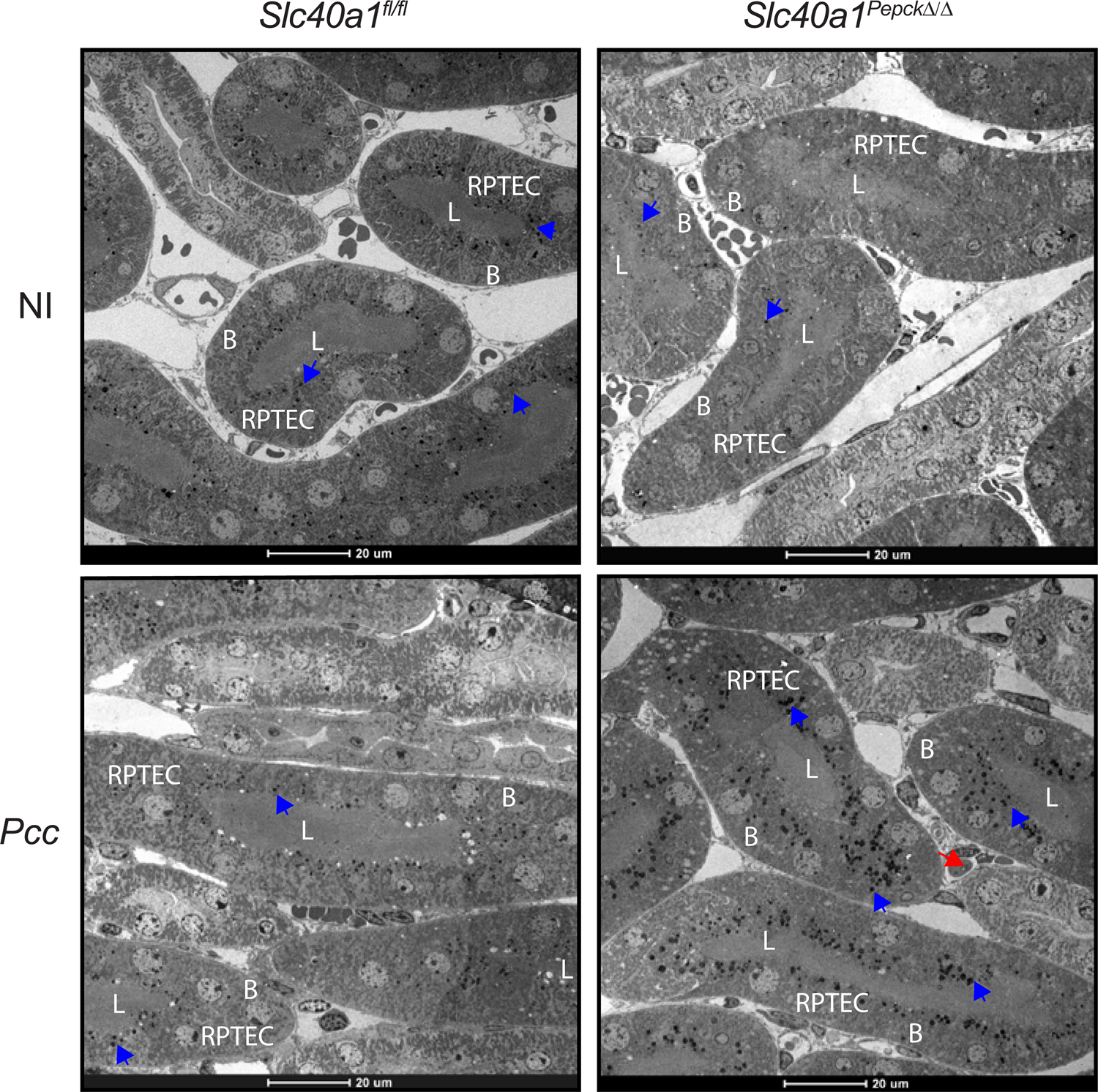
Impact of Slc40a1 expression on RPTEC damage. Electron microscope images from the renal cortex of non-infected (NI) and at the peak of *Pcc* infection (Day 7) in *Slc40a1^Pepck^*^Δ/Δ^ and *Slc40a1*^fl/fl^ mice. Images are representative of 2 animals *per* genotype and experimental condition. Blue arrows indicate electron-dense deposits, most likely corresponding to Fe. Red arrows highlight infected RBC. L: renal proximal tubule lumen; B: RPTEC basolateral surface. Scale bars: 20µm.

**Extended Data Figure 16.**
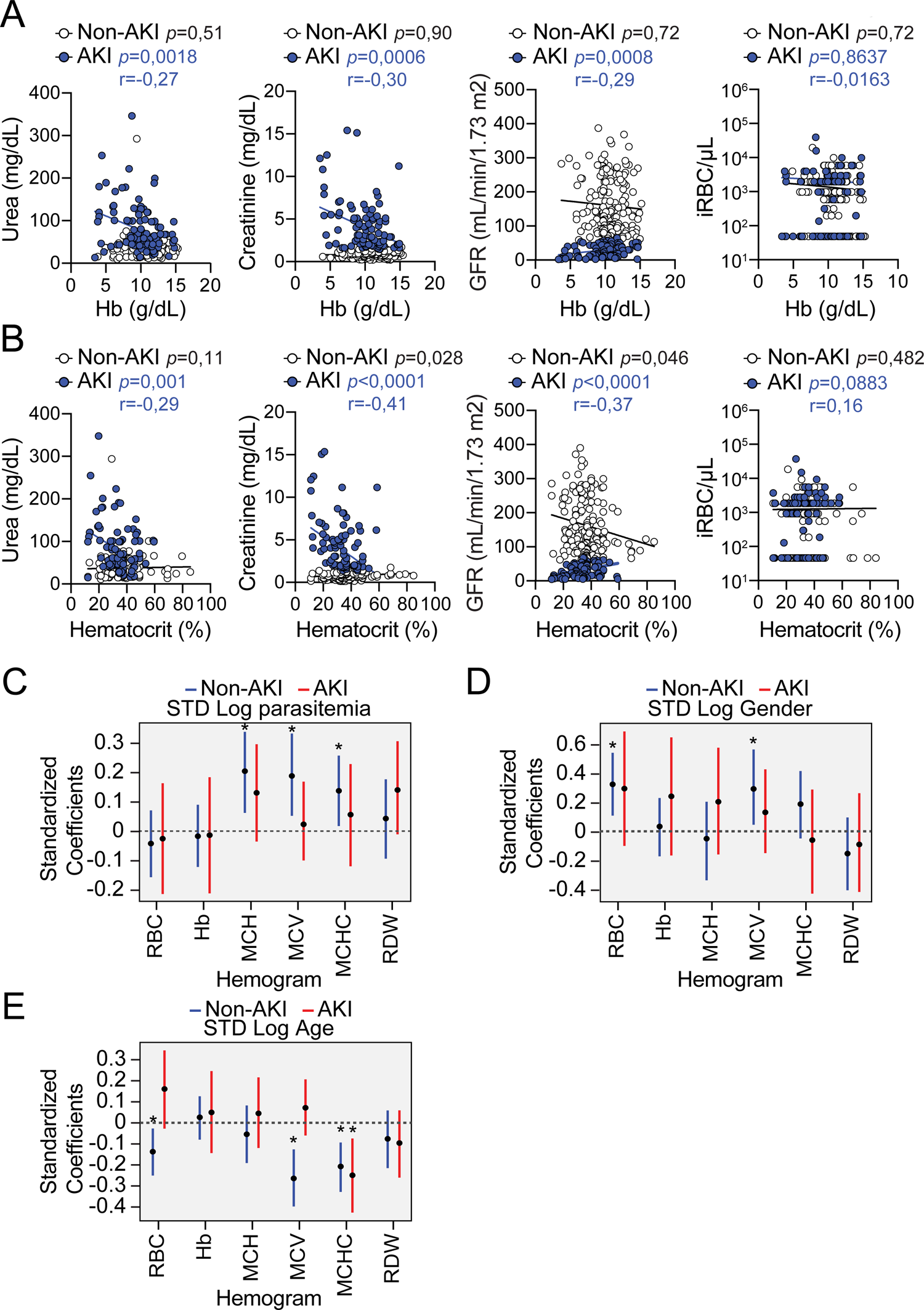
AKI predicts anemia severity in *P. falciparum*-infected individuals. **A, B**) Correlation coefficients between serum urea, creatinine, glomerular filtration rate (GFR) and parasitemia (iRBC/µL), and: **A)** hemoglobin concentration or **B)** hematocrit in *P. falciparum*-infected patients with acute kidney injury (AKI) or without AKI (non-AKI). For each correlation the corresponding *p* value for non-AKI (black) or AKI (blue) patients is highlighted. Spearman’s correlation coefficient (r) is indicated for AKI (blue) patients. Same patients as in (*Fig. 4H*). **C-E)** Quantification of associations between **C)** parasitemia, **D)** age and **E)** gender and hemogram outputs after adjustment for gender, age and AKI status. Each bar depicts the estimate and uncertainty of the standardized coefficient (95% HPD) for age or sex, of a linear regression with the indicated hemogram output as response variable and urea, parasitemia, AKI status, age and gender as independent variables. Same patients as in (*Fig. 4H*).

**Figure S_1.**
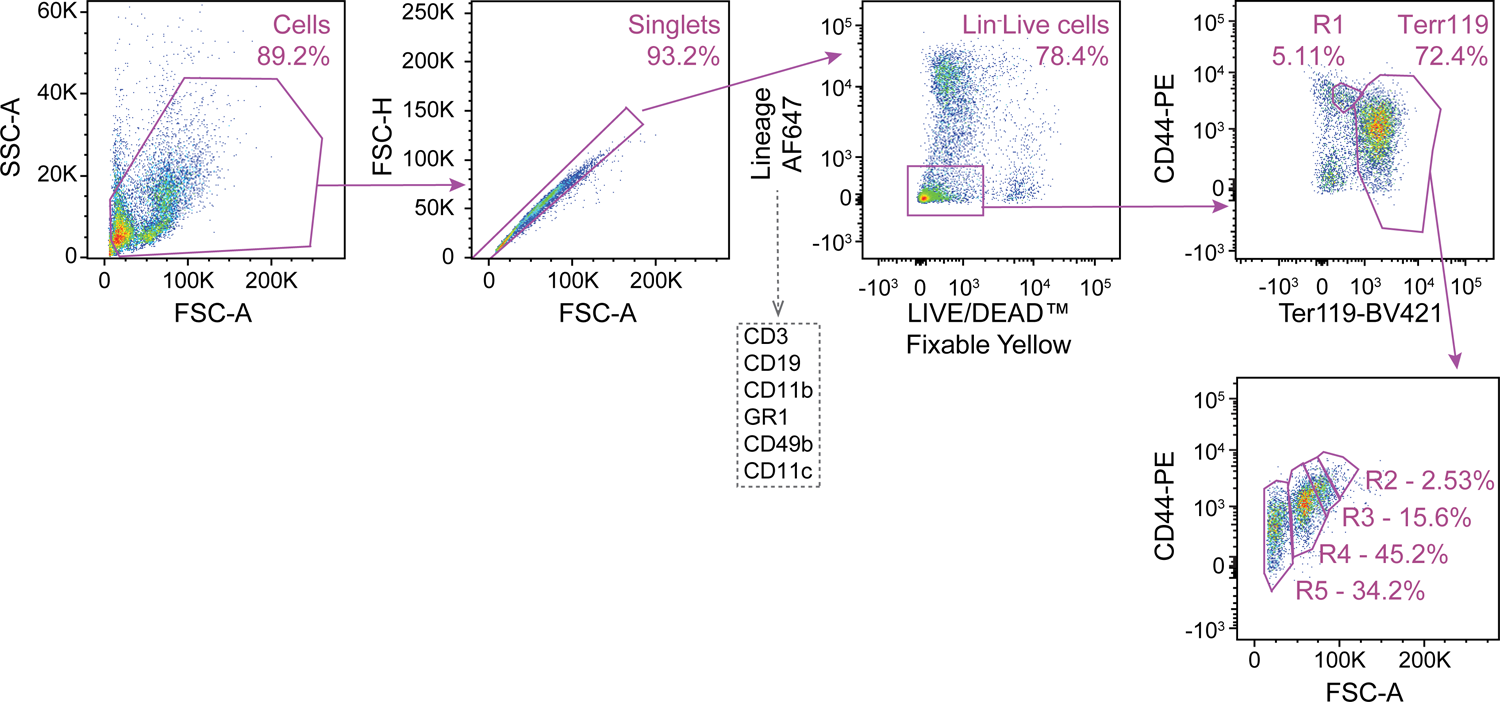
Gating strategy for the identification of RBC developmental stages. Spleen and bone marrow cells were plotted according to size (FSC-A) *vs.* granularity (SSC-A). Cell doublets were excluded according to relative area (FSC-A) *vs.* height (FSC-H) ratio, selecting single “round” cells. Dead cells were excluded based on negative staining for the LIVE/DEAD™ Fixable Yellow Dead Cell Stain. Non-erythroid cells were excluded according to the expression of CD3, CD19, CD11b, GR1, CD49b or CD11c lineage markers. Proerythroblasts (R1) were identified as CD44^high^Ter119^int^ cells. Ter119^high^ cells were segregated into basophilic erythroblasts (R2), polychromatic erythroblasts (R3), orthochromatic erythroblasts and reticulocytes (R4) and mature RBC (R5) according to CD44 expression levels and relative size (FSC-A), as described^42^.

**Figure S_2.**
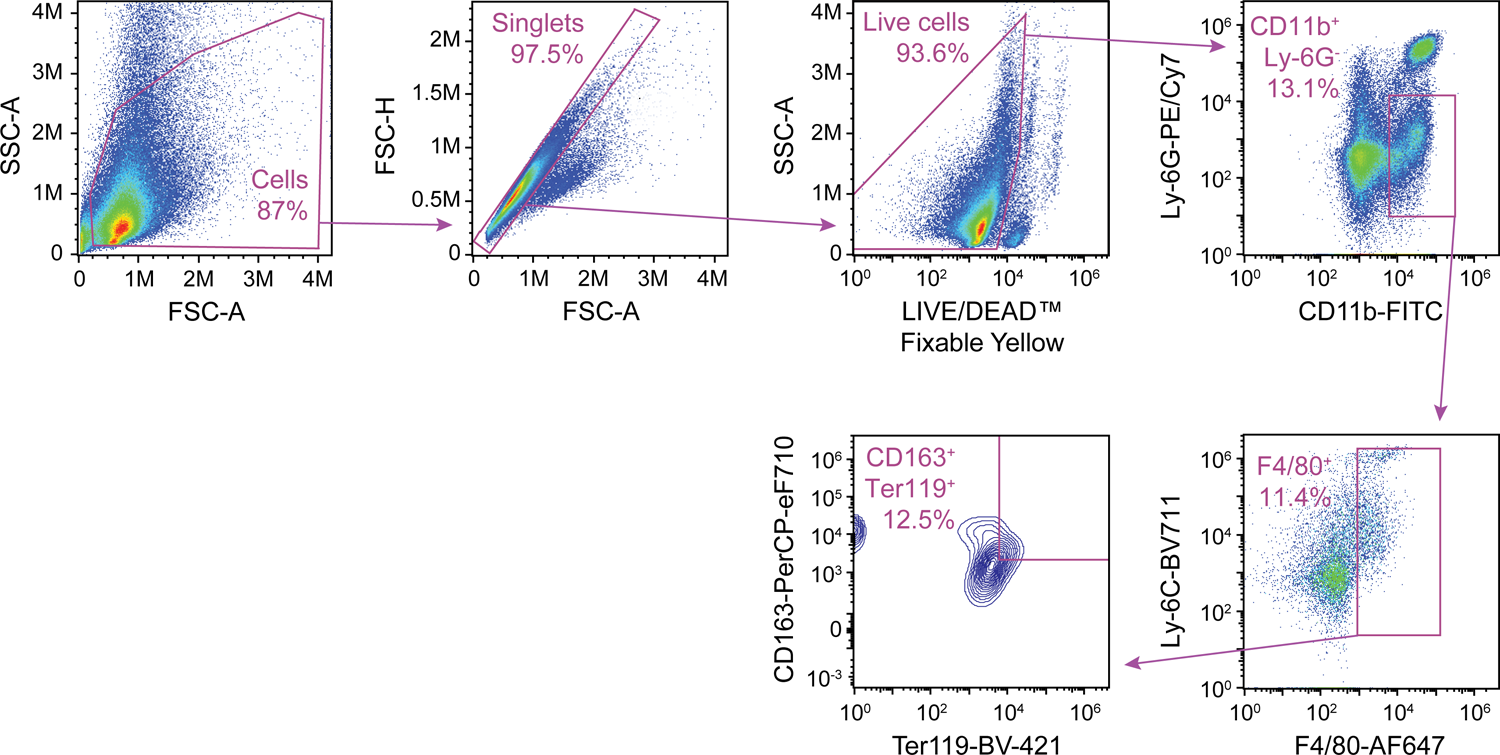
Gating strategy for the identification of erythrophagocytic macrophages. Leukocytes were identified in size (FSC-A) *vs.* granularity (SSC-A) plots. Cell doublets were excluded according to cellular area (FSC-A) *vs.* height (FSC-H), selecting for single “round” cells. Live cells were selected based on negative staining for the LIVE/DEAD™ Fixable Yellow Dead Cell Stain and myeloid cells, excluding granulocytes, were selected as CD11b^+^Ly-6G^-^ cells. Erythrophagocytic macrophages were identified based on positive staining for Ly-6C, F4/80, CD163 and Ter119 (i.e., CD11b^+^Ly-6C^int^Ly-6G^-^F4/80^+^CD163^+^Ter119^+^).

**Figure S_3.**
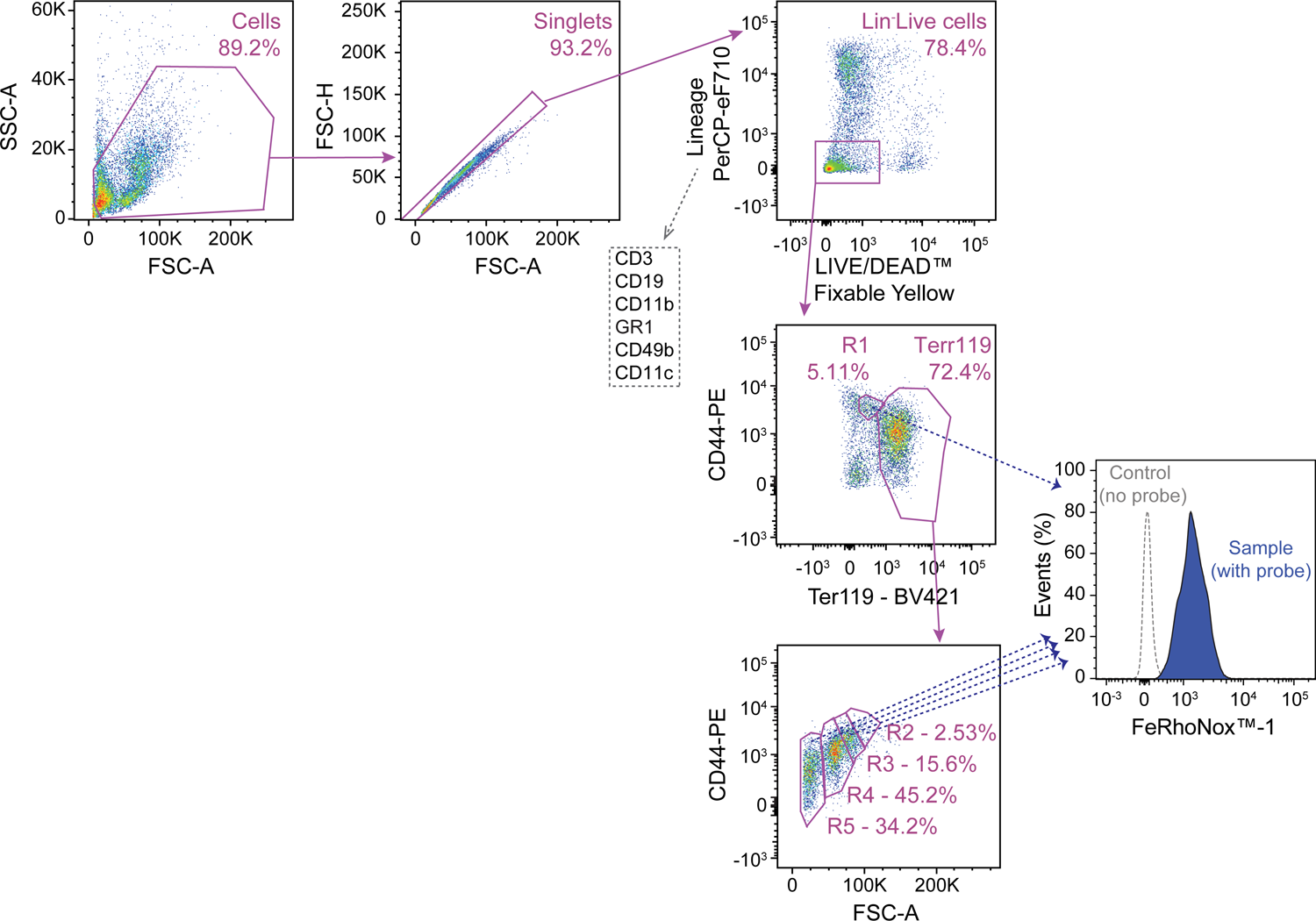
Gating strategy for erythroblast Fe^2+^ staining. Erythroblasts were identified using the same gating strategy as in *Figure S_1*. Relative intracellular Fe^2+^ quantification was assessed by evaluating FeRhoNox^TM^-1 staining in each erythroblast population (*i.e.,* R1-R5).

## Methods

### Mice

Mice were bred and maintained under specific pathogen-free (SPF) conditions at the Instituto Gulbenkian de Ciência (IGC). Protocols were approved in a two-step procedure, by the Animal Welfare Body of the IGC and by the Portuguese National Entity that regulates the use of laboratory animals in research (DGAV - Direção Geral de Alimentação e Veterinária). Experiments on mice followed the Portuguese (Decreto-Lei n° 113/2013) and European (Directive 2010/63/EU) legislation, concerning housing, husbandry and animal welfare. C57BL/6J wild-type mice were purchased from the IGC animal facility. C57BL/6J *Pepck*^Cre^ mice were previously described^17, 43^. For the generation of C57BL/6J *Slc40a1* floxed (*Slc40a1^fl/fl^*) mice, sperm was obtained from EMMA repository (EM:04833) and revitalized in C57BL/6J oocytes (IGC, Transgenesis facility). *Slc40a1^fl/WT^* mice were crossed with C57BL/6J *Flp* recombinase mice (Jackson Laboratory; #009086) to remove the floxed neomycin/LacZ cassette (*Fig. S4A*). For genotyping, DNA was extracted from a small earpiece and amplified by PCR (Xpert directXtract PCR Kit, GRiSP, #GE60.0480) using the following primer sets: SLC40A1-5arm-WTF: AAACAGCAAAGACTTAAAAGATGGA, SLC40A1-Crit-WTR: GTTCACTGCACCAG CATGTC, 5mut-R1: GAACTTCGGAATAGGAACTTCG, 5’CAS-F1:AAGGCGCA TAACGATACCAC, 3’CAS-R1:CCGCCTACTGCGACTATAGAGA, as described in: https://www.infrafrontier.eu/*; Slc40a1 HEPD0551_2_A0*7. PCR products were resolved (2% agarose; GRS Ladder 100bp; GRiSP, #GL041.0050)(*Fig. S4A*). *Pepck*^Cre/Wt^ mice were crossed with *Slc40a1^fl/fl^* mice to generate *Pepck*^Cre^*Slc40a1*^fl/fl^ (*Slc40a1^Pepck^*^fl/fl^) and littermate *Slc40a1^fl/fl^* mice (*Fig. S4A*). Conditional deletion of the *Slc40a1^fl/fl^* allele in RPTEC from *Slc40a1^Pepck^*^Δ/Δ^ mice was induced 8-week after birth by the addition of 0,3M NH_4_Cl to drinking water during one week, as described^17, 43^ (*Fig. S4A-D*). *Slc40a1^Pepck^*^Δ/Δ^ mice were used for experimental procedures 4 weeks after exposure to acidified water. The *Egfp-L10^fl/Wt^* (EF1a-Flox-GFP-L10; Kind gift from Dr. Ana Domingos, Oxford University, UK)^19^ were crossed with *Pepck*^Cre/Wt^ and *Pepck^Cre/Wt^Slc40a1^fl/fl^*, to generate *Egfp-L10^Pepck^* (*Fig. S2A*) and *Slc40a1^Pepckfl/fl^Egfp-L10^Pepck^* (*Fig. S4E-F*) mice, respectively. Deletion of the *Slc40a1^fl/fl^* allele in RPTEC from *Slc40a1^PepckΔ/Δ^Egfp-L10^Pepck^* was induced as described above (*Fig. S4E-F*).

### RPTEC isolation and treatment

C57BL/6 mice were sacrificed and perfused *in toto* (PBS, 10mL). The kidneys were harvested and the renal cortex dissected from the medulla, minced finely and passed through a 70 µm cell strainer in RPTEC media (Renal Epithelial Cell Growth Medium 2, PromoCell, # C-26130) supplemented with Penicillin-Streptomycin Solution (100U/mL, Biowest, L0022-100) and supplements provided in the RPTEC media (5% Fetal Calf Serum, 10ng/mL Epidermal Growth Factor (recombinant human), 5µg/mL Insulin (recombinant human), 0.5µg/mL Epinephrine, 36ng/mL Hydrocortisone, 5µg/mL Transferrin (recombinant human), 4pg/mL Triiodo-L-thyronine). Tissue homogenates were centrifuged (154*g,* 5min., RT) and pellets washed (2x in RPTEC media). Cell suspensions were plated on collagen-coated 10cm dishes (NeoTC Cell Culture Dish 100×20mm, standard growth surface for adherent cells, Sarstedt, #83.3902) and incubated (ON, 37°C, 5% CO_2_). Media containing renal tubules was collected, centrifuged (154*g*, 5min. RT) and tubules were resuspended in RPTEC media, plated on collagen-coated 12-well plates and incubated (37°C, 5% CO_2_), replacing media every third day. At confluency (∼7 days) RPTEC were exposed to hemin (10µM, Frontier Scientific, #FSIH651) in serum-free RPTEC media for the indicated times.

### *Plasmodium* infection and disease assessment

Mice were infected with *Plasmodium chabaudi chabaudi* AS (*Pc*AS) or transgenic GFP-expressing *Pc*AS (*i.e.,* PcAS-GFP_ML_)^44^ by intra-peritoneal (i.p.) inoculation of fresh blood collected from infected mice (2×10^6^ infected RBC; iRBC). *P. chabaudi chabaudi* infection is referred to, throughout the text, as *Pcc*. Mice were monitored daily for parasitemia, body weight, core body temperature, RBC number and survival. The day of infection was considered as zero (D0). Rectal temperature was determined using a Rodent Thermometer BIO-TK8851 (Bioset). Body weight was monitored using an Ohaus^®^ CS200 scale (Sigma Aldrich). Number of RBC per mL of blood was quantified by flow cytometry on a LSR Fortessa X20 analyzer (BD Biosciences) using a known concentration of reference 10µm latex beads suspension (Coulter® CC Size Standard L10, Beckman Coulter, # 6602796), gating on RBC in blood samples, based on size and granularity and on bead population. Percentage of *Pcc*AS iRBCs (i.e., parasitemia) was determined manually by optical microscopy, counting *Plasmodium* containing RBC in 4 fields of Giemsa-stained blood smears (1000x magnification). PcAS-GFP_M_ parasitemia was determined by flow cytometry, according to the percentage of GFP^+^ RBC. Pathogen load (parasitemia x RBC number) was expressed as iRBC/µL. Disease trajectories were represented by plotting the median values of each disease parameter against the median pathogen load during infection, as described^7^.

### Fe quantification

Non-heme Fe quantification was performed in mouse tissues, as described^7^. Briefly, tissues were harvested, weighted, digested (1mL; 3M HCL, 0.61M trichloroacetic acid, 50h, 65°C), routinely vortexed during digestion and centrifuged (12,000*g,* 1min., RT). Non-heme Fe was detected in the supernatants using a chromogenic assay, whereby the chromogen reagent solution was prepared fresh using 1 volume of the chromogen stock solution (1.86mM bathophenanthroline sulfonate-Sigma, #B-1375, 143mM thioglycolic acid-Sigma, #T-6750 in milliQ water), 5 volumes of saturated sodium acetate (Sigma, #S5636) and 5 volumes of ion-free water. Samples (10µL) were incubated (15min., RT) with the chromogen reagent solution (200µL) in 96-well plates and optical density (λ_535_ nm) was measured in a Multiskan Sky microplate reader (Thermo Scientific). Ferrous chloride (10µL; 500µg/dL) was used as standard. Non-heme Fe was expressed in µg *per* tissue or *per* g of wet weight tissue.

### Lipid peroxidation

Malondialdehyde (MDA) was quantified using Thiobarbituric Acid Reactive Substances (TBARS, TCA Method, Cayman, cat #700870). Briefly, mice were sacrificed and perfused (PBS, 10mL, RT), tissues collected, sliced on ice, weighted, homogenized in RIPA buffer (150mM NaCl, 1% Nonident P-40, 0.5% DOC, 0.1% SDS in Tris 50mM, pH 7.4) containing protease inhibitors (cOmplete™, Mini, EDTA-free Protease Inhibitor Cocktail, Roche #11836170001) and centrifuged (1600g, 10min.; 4°C). Supernatants (100µL) were mixed with TCA Assay Reagent (10%, 100µL) and color reagent (800µL), vortexed, heated (95°C, 1h) and immediately placed on ice (10min.). Samples were centrifuged (1,600*g*, 10min., 4°C) and supernatant absorbance was read (λ_540_ nm) in a Multiskan Sky microplate reader (Thermo Scientific). Serial dilutions of purified MDA were used as a standard and MDA concentration was expressed as µmol MDA/g wet weight tissue.

### Serum Fe and NTBI

Fe concentration and transferrin saturation in serum were quantified using the Iron/TIBC Reagent Set, according to manufacturer’s instructions (Pointe Scientific, #1750460). Non-transferrin bound iron (NTBI) concentration in serum was quantified using a nitrilotriacetic acid (NTA) assay, as described^45^. Briefly, plasma was mixed with NTA (Sigma, #N0128 and #N0253, in 1:1 ratio) solution (800mM; 9 parts plasma: 1 part NTA), incubated (30min., RT) and centrifuged (3000*g*, 1h, 4°C) in an Ultra-2 Centrifugal Filter Unite (Millipore, #UFC203024). Filtered samples (100µL) were incubated (1h; RT) with MOPS buffer (5mM, 100µL, Sigma, #M1254), Bathophenanthrolinedisulfonic acid disodium salt hydrate (BPT, 60mM, 25µL, Sigma, #B1375) and Thioglycolic acid solution (TGA, 120mM, 25µL, Sigma, #T6750). Serial dilutions of ammonium iron (III) sulfate dodecahydrate (Sigma, #221260) were used as standard. Absorbance was measured (λ_537_ nm) in a Multiskan Sky microplate reader (Thermo Scientific).

### Hemogram and Serology

Mice were sacrificed and blood was obtained at the indicated time-points after *Plasmodium* infection. Complete blood counts (hemograms) as well as urea, creatinine, aspartate aminotransferase (AST), alanine aminotransferase (ALT), lactate dehydrogenase (LDH), Transferrin and Troponin I plasma concentrations were measured by DNATech (Portugal; http://www.dnatech.pt/web/).

### Histology

Mice were perfused *in toto* with PBS (1X, 10mL) and organs were harvested, fixed (10% formalin), embedded in paraffin, sectioned (3µm) and stained with Hematoxylin & Eosin (H&E) or with Perl’s Prussian Blue, as described^46^. Whole sections were analyzed in a DMLB2 microscope (Leica), images were acquired with a DFC320 camera (Leica) and a NanoZoomer-SQ Digital slide scanner (Hamamatsu Photonics). Images were reconstructed using the NDP.view2 (Hamamatsu Photonics) software. Histopathology analyzes were performed by Dr. Pedro Faísca (IGC Histopathology Unit). For iron (Fe^3+^) quantification in kidney sections, the fraction of iron positive staining the total area of the kidney was quantified using the color threshold plugin of the Image J software (Rasband, W.S., ImageJ, U.S. NIH, Bethesda, Maryland, USA).

### Electron Microscopy

Mice were sacrificed, perfused *in toto* (PBS; 10mL) and fixed (2% formaldehyde EMS #15710, 2.5% glutaraldehyde EMS #16220 in 0.1M Phosphate Buffer). Kidneys were harvested and fixed in the same solution, dissected, and the junctional region between cortex and medulla (to assure sampling of both regions in the same section) was cut into small pieces (< 2mm) and immersed in fixative (overnight, 4°C). Samples were microwave-fixed (PELCO Biowave Pro+, Ted Pella, 7 cycles, 2min. each, alternating irradiation power of 100w and 0w) washed (3x in 0.1M phosphate buffer) and incubated with 1% osmium tetroxide (EMS #19110) in 0.1 phosphate buffer (8 cycles as in initial fixation). Samples were washed (2x in 0.1M phosphate buffer, 2x in dH_2_O), stained with 1% Tannic acid (EMS #21700), washed (5x 5min. in dH_2_O) and incubated with 1% UA (Analar #10288) in dH_2_O. The staining steps were done in the microwave (7x 1min. cycles) with alternating irradiation power (150 and 0w). Samples were dehydrated in a graduated ethanol series (30%, 50%, 75%, 90% and 100% 3x) using a cycle of 40sec. each with 150W power and infiltrated in EMbed-812 epoxy resin (EMS #14120) with incremental resin concentrations (25%, 50%, 75%, 100%), before polymerization (ON, 60°C). Sections (70nm) of were cut (UC7 Ultramicrotome, Leica) and stained with uranyl acetate and lead citrate (5min. each) before analyzes in a Tecnai G^2^ Spirit BioTWIN Transmission Electron Microscope (TEM) from FEI operating at 120 keV and equipped with an Olympus-SIS Veleta CCD Camera. Image analysis was performed using ImageJ (Rasband, W.S., ImageJ, U.S. NIH, Bethesda, Maryland, USA).

### Mouse Metabolic Monitoring

The Promethion behavioral and phenotyping system used consists of a standard GM-500 cage with a food hopper and a water bottle connected to load cells (2 mg precision) with 1 Hz rate data collection. Each cage contains red house enrichment. Ambulatory activity was monitored at 1 Hz rate using an XY beam break array (1 cm spacing). Oxygen, carbon dioxide and water vapor were measured using a CGF unit (Sable Systems). The multiplexed system was operated in pull-mode. Air flow was measured and controlled by the CGF (Sable Systems) with a set flow rate of 2 L/min. O_2_ consumption and CO_2_ production were reported in milliliters per minute (mL/min). Energy expenditure was calculated using the Weir equation^47^ and Respiratory Exchange Ratio (RER) was calculated as the ratio of VCO_2_/VO_2_. Raw data was processed using Macro Interpreter v2.41 (Sable Systems). *Slc40a1^Pepck^*^Δ/Δ^ and *Slc40a1^fl/fl^* mice were housed on a 14/10 h light/dark cycle, acclimatized for two days and analyzed thereafter at steady state (*i.e.,* not infected). The same mice were infected with *Pcc* and infection was allowed to proceed in regular housing. Five days post-infection, mice were re-housed in the Promethion system to resume recordings for additional 5 days (*i.e.,* 5-10 post-infection).

### Translating Ribosome Affinity Purification (TRAP)

was adapted from Dr. McMahon (University of South California, USA)^48^ and Dr. N. Heintz (Rockefeller University, NYC, USA)^49^.

#### Affinity matrix preparation

Streptavidin MyOne T1 Dynabeads (150µL *per* kidney, Thermo fisher, #65602) were washed (1x in PBS) and collected on magnet (>1min.), discarding the supernatant. Beads were re-suspended in PBS (500µL, 1X, Thermofisher #AM9625), diluted in RNAse-free water (Thermofisher #AM9939) and incubated with Pierce^TM^ Recombinant Protein L Biotinylated (60µL *per* kidney, ThermoFisher #29997) in a tube rotator (25min., RT). Beads were collected on magnet (>1min.), discarding the supernatant, washed (4x in PBS, 3% BSA) and re-suspended (500µL *per* kidney) in low salt buffer (20mM HEPES-KOH, Sigma Aldrich #H0527, 0,15M KCl, Thermofisher #AM9640G, 10mM MgCl2, Thermofisher #AM9530G, 1% NP-40, Hölzel Diagnostika Handels GmbH #P-1505, 100µg/mL Cycloheximide, Sigma #C7698, 0,5mM Dithiothreitol, DTT, Sigma, #D9779, in RNAse-free water). Beads in low salt buffer were incubated (1h, 4°C) in a tube rotator with monoclonal anti-GFP antibodies (25µg, Core Facility, Memorial Sloan-Kettering Cancer Center, New York, USA, clone numbers 19F7 and 19C8). Beads were collected on magnet (>1min.) discarding the supernatant, washed (3x) and re-suspended in low salt buffer (200µL).

#### Tissue homogenates preparation

*Egfp-L10^Pepck^* and *Slc40a1^PepckΔ/Δ^Egfp-L10^Pepck^* mice were sacrificed, perfused (10mL, ice-cold PBS) and kidneys were harvested, minced in ice-cold dissection buffer (2mL *per* kidney, 1X HBSS, Thermofisher #14065056, 2,5mM HEPES-KOH Sigma Aldrich #H0527, 35mM D-Glucose, Sigma #G7528, 4mM M sodium bicarbonate, Sial #S6297, 100µg/mL Cycloheximide, Sigma #C7698, in RNAse-free). Kidney slices were homogenized in lysis buffer (1mL, 20mM HEPES-KOH Sigma Aldrich #H0527, 0,15M KCl 10mM #AM9640G, 10mM MgCl_2_ Thermofisher #AM9530G, 100µg/mL Cycloheximide, Sigma #C7698, in RNAse-free water), supplemented with 1 tablet of Protease inhibitors (cOmplete™, Mini, EDTA-free Protease Inhibitor Cocktail, Roche #11836170001) *per* 10ml lysis buffer, and 10µL RNasin RNase inhibitor (PROMN2515, Promega) and 10µl Superasin RNase inhibitor (AM2696, Thermofisher) *per* mL lysis buffer. Tissue lysates were centrifuged (2,000*g*, 10min., 4°C) and supernatants were supplemented with NP-40 (1%, Hölzel Diagnostika Handels GmbH #P-1505) and 1,2-diheptanoyl-sn-glycero-3-phosphocholine (DHPC, 30mM, Avanti Polar Lipids ##850306P), incubated (5min., on ice) and centrifuged (20,000*g*, 15min., 4°C). Supernatants were collected for the next steps.

#### Immunoprecipitation

Kidney lysates (1mL) were incubated with affinity matrix beads (200µL; 30min. 4°C). The mixtures were placed on pre-chilled magnets (1min.) and the supernatant was discarded. Beads were washed with High salt buffer (50mL, 20mM HEPES-KOH Sigma Aldrich #H0527, 0,35M KCL Thermofisher #AM9640G, 10mM MgCl_2_ Thermofisher #AM9530G, 1% NP-40 Hölzel Diagnostika Handels GmbH # P-1505, 100µg/mL Cycloheximide, Sigma #C7698, 0,5mM Dithiothreitol, DTT, Sigma, #D9779, in RNAse-free water). Beads were removed from the magnet, incubated (5min., RT), resuspended in RNA extract lysis buffer (100µL RLT lysis buffer from RNA extraction kit supplemented with 40mM DTT, Sigma, #D9779), vortexed and let sit (10min., RT) to allow for RNA to be released. RNA was extracted from the beads using RNeasy Micro Kit (QIAGEN, #74004), as *per* manufacturer’s instructions.

### Bulk RNA sequencing and TRAP RNA Analysis

Extracted RNA was assessed for quality using a 2100 Bioanalyzer (Agilent Technologies, #5067-1513) in combination with the RNA 6000 pico kit (Agilent Technologies). Full-length cDNAs and sequencing libraries were generated according to the SMART-Seq2 protocol, as described^50^. Following quality control, library preparation including cDNA ‘tagmentation’, PCR-mediated adaptor addition and amplification of libraries was performed using the Nextera library preparation protocol (Nextera XT DNA Library Preparation kit, Illumina), as described^51^. Libraries were sequenced (NextSeq 500, Illumina) using High Output kit v2.5 (75 cycles). Sequence information was extracted in FastQ format, using Illumina’s bcl2fastq v.2.19.1.403, producing on average 32.54×10^6^ reads *per* sample. Library preparation and next-generation sequencing were performed at the IGC Genomics Unit.

Fastq reads were aligned against the mouse reference genome GRCm39 using the GENCODE vM27 annotation to extract splice junction information (STAR; v.2.5.2a)^52^. Read summarization was performed by assigning uniquely mapped reads to genomic features using *FeatureCounts* (v.1.5.0-p1). Gene expression tables were imported into the R programming language and environment (v.4.1.0) to perform differential gene expression and functional enrichment analyses, as well as data visualization.

Differential gene expression was performed using the DESeq2 R package (v.1.32). Gene expression was modeled by genotype and condition, which included the following factors: *Slc40a1^fl/fl^* (control) or *Slc40a1^Pepck^*^Δ/Δ^ mice, which were either non-infected (NI) or infected with *Pcc* (N=3 for each combination). Genes not expressed or with fewer than 10 counts across the 12 samples were removed, leaving 19,052 genes for downstream differential gene expression analysis. We subsequently ran the function *DESeq* to estimate the size factors (by *estimateSizeFactors*), dispersion (by *estimateDispersions*) and fit a binomial GLM fitting for βi coefficient and Wald statistics (by *nbinomWaldTest*). Pairwise comparisons tested with the function *results* (alpha = 0.05), were: 1) *Pcc*-infected *Slc40a1^fl/fl^ vs.* NI *Slc40a1^fl/fl^*; 2) *Pcc*-infected *Slc40a1^Pepck^*^Δ/Δ^ vs. *Pcc-infected Slc40a1^fl/fl^*; 3) NI *Slc40a1^Pepck^*^Δ/Δ^ vs. NI *Slc40a1^fl/f^* and 4) *Pcc*-infected *Slc40a1^Pepck^*^Δ/Δ^ *vs.* NI *Slc40a1^Pepck^*^Δ/Δ^. In addition, the log_2_ fold change for each pairwise comparison was shrunken with the function *lfcShrink* using the algorithm *ashr* (v.2.2-47)^53^. Differentially expressed genes were considered for genes with an adjusted p-value<0.05 and an absolute log_2_ fold change>0. Normalized gene expression counts were obtained with the function *counts* using the option normalized = TRUE. Regularized log transformed gene expression counts were obtained with *rlog*, using the option blind = TRUE. Principal Component Analysis (PCA) of overall sample expression profiles was performed with function *pcaPlot* from the DESeq2 R package (v.1.32), using regularized log transformed gene expression counts for each sample and grouped according to “condition”. Probability ellipses were calculated using a function adapted from *pcaplot* from the Bioconductor pcaExplorer R package (v.2.18)^54^.

Ensembl gene ids were converted into gene symbols from Ensembl (v.104 - May 2021-https://may2021.archive.ensembl.org) by using the mouse reference (GRCm39) database with biomaRt R package (v.2.48.2). All scatter plots, including volcano plots, were done with the ggplot2 R package (v.3.3.5). Heatmaps were made with pHeatmap (v.1.0.12), using the Euclidean distance and Ward.D2 method for clustering estimation. For hierarchical clustering of differentially expressed genes, gene expression counts were scaled (Z-score) with the function *scale*.

Functional enrichment analysis was performed with the gprofiler2 R package (v.0.2.1). Enrichment was performed using the function *gost* based on the list of up- or down-regulated genes (genes with an adjusted p-value<0.05 and a log_2_ fold-change>0 or <0), between each pairwise comparison (independently), against annotated genes (domain_scope = “annotated”) of the organism *Mus musculus* (organism = “mmusculus”). Gene lists were sorted according to adjusted p-value (ordered_query = TRUE) to generate GSEA (Gene Set Enrichment Analysis) style *p*-values. Only statistically significant (user_threshold=0.05) enriched functions are returned (significant=TRUE) after multiple testing corrections with the default method g:SCS (correction_method = “analytical”). The gprofiler2 queries were run against all the default functional databases for mouse which include: Gene Ontology (GO:MF, GO:BP, GO:CC), KEGG (KEGG), Reactome (REAC), TRANSFAC (TF), miRTarBase (MIRNA), Human phenotype ontology (HP), WikiPathways (WP), and CORUM (CORUM). For future reference, gprofiler2 was performed using database versions Ensembl 104, Ensembl gene 51 (database updated on 07/05/2021). For STRING database network analysis, genes contained within enriched gene sets associated with Type I and II interferon responses, Fe homeostasis, Glutathione biosynthesis and oxidative stress response were merged and uploaded to the STRING database (v11.5) and queried for known protein-protein interactions (organism: *Mus musculus*; interaction score > 0.4). The resulting network was imported into Cytoscape (v3.9.0) for network layout design. RNA sequencing data was deposited at the NCBI GEO with the accession number GSE189579.

### qRT-PCR

Mice were sacrificed, perfused (10mL PBS), organs were harvested, snap-frozen in liquid nitrogen and RNA was extracted using tripleXtractor reagent (GRISP, #GB23.0100). Total RNA was used for cDNA synthesis (GRISP, # GK81.0100), followed by qPCR using Power SYBR Green PCR master mix (Bio-Rad, #1725124) on an ABI QuantStudio^TM^ 7 Flex system (Thermo Scientific). Transcript values were calculated from the threshold cycle (Ct) of each gene using the 2^-ΔΔCT^ method and normalized to Acidic ribosomal phosphoprotein P0 (*Arbp0*) or beta-Actin (*Actin*). Primers for qPCR include: *Arbp0*, Fwd-CTTTGGGCATCACCACGAA, Rev-GCTGGCTCCCACCTTGTCT; *Slc40a1*, Fwd-TGCCAGACTTAAAGTGGCCC, Rev-GCAGACAGTAAGGACCCATCC; *Lcn2*, Fwd-GCCCAGGACTCAACTCAGAA, Rev-GACCAGGATGGAGGTGACAT; *Kim1*, Fwd-GGAAGTAAAGGGGGTAGTGGG, Rev-AAGCAGAAGATGGGCATTGC; *Epo*, Fwd-TGGTCTACGTAGCCTCACTTCACT, Rev-TGGAGGCGACATCAATTCCT; *Erfe*, Fwd-ATGGGGCTGGAGAACAGC, Rev-TGGCATTGTCCAAGAAGACA; *EpoR*, Fwd-GGACCCTCTCATCTTGACGC, Rev-CTTGGGATGCCAGGCCAGAT; *Hamp1*, Fwd-GAGAGACACCAACTTCCCCA, Rev-TCAGGATGTGGCTCTAGGCT. *Tfr1*, Fwd-GTTTTTGTGAGGATGCAGACTATCC, Rev-GCTGAGGAACTTTCTGAGTCAATG. *Actin*, Fwd-AAATCGTGCGTGACATCAAAGA, Rev-GCCATCTCCTGCTCGAAGTC.

### Western Blotting

Tissue or cells were lysed in 2% SDS-PAGE sample buffer (100mM Tris, pH 6.8, 20% glycerol, 4% SDS, 0.002% bromophenol blue, 100mM DTT, protease inhibitor cocktail: Sigma, #P8340). Total protein was quantified at λ_280_nm (Nanodrop 2000; ThermoFisher Scientific), resolved (50µg) on a 12% SDS-PAGE and transferred to Polyvinylidene fluoride (PVDF) membranes. These were blocked (5% skim milk in TBS-T), washed (1x in TBS-T) and incubated with primary antibodies (Overnight, 4°C). Primary antibodies included: Rabbit polyclonal anti-Slc40a1 (1:1000)^55^, goat polyclonal anti-Gapdh (Sicgen, #AB0049-200, RRID: AB_2333141; 1:4000). Membranes were washed (1x in TBS-T) and incubated (1h, RT) with peroxidase-conjugated secondary antibodies: AffiniPure Goat Anti-Rabbit IgG (H+L; polyclonal)(Jackson Immunoresearch, #111-035-045; RRID: AB_2337938; 1:5000), AffiniPure Goat Anti-Mouse IgG (H+L; polyclonal)(Jackson Immunoresearch, #111-035-062; RRID: AB_2338504; 1:5000), or donkey anti-Goat IgG polyclonal antibody (ThermoFisher Scientific, #PA1-28664, RRID: AB_10990162, 1:5000). Membranes were washed (1x in TBS-T) and peroxidase activity was detected using SuperSignal™ West Pico PLUS Chemiluminescent Substrate (ThermoFisher Scientific, cat# 34580). Blots were developed using Amersham Imager 680 (GE Healthcare), equipped with a Peltier cooled Fujifilm Super CCD. Densitometry analysis was performed with ImageJ (Rasband, W.S., ImageJ, U.S. NIH, Bethesda, Maryland, USA, https://imagej.nih.gov/ij/,1997-2014), using only images without saturated pixels.

### Immunofluorescence and super-resolution imaging of kidney cryosections

Co-staining of Slc40a1 in the kidney of *Egfp-L10*^Pepck^ mice, expressing EGFP specifically in RPTEC (*Fig. S2A*) was performed essentially as described^56^. Briefly, mice were sacrificed, perfused (10mL; ice-cold PBS) and kidneys were immediately harvested, fixed (4% paraformaldehyde; Merck, #1.04005.1000 in PBS)(24h; 4°C), washed (30min. in PBS, 2x) and soaked in 12.5% (2h; 4°C) and 25% (7 days, 4°C) sucrose (Sigma). Kidneys were embedded in OCT (Tissue-Tek®, SAKURA, R1180), frozen in dry ice and sectioned (10µm; Leica Cryostat CM 3050 S; Leica Biosystems). Sections were soaked (2 x 5min.) in PBS, 0.02% Tween20 (VWR, hereafter PBS-T), permeabilized (2 x 10min.) in PBS, 0.2% Triton X-100 (Sigma Aldrich), washed in PBS (2x; 10min), blocked (3% BSA in PBS-T, 2h) and incubated with a rabbit polyclonal anti-Slc40a1^55^ (1:250, in 3% BSA in PBS-T, overnight, 4°C) in a humidified chamber. Negative controls were performed by omitting the primary antibody. Sections were washed in PBS-T (4x 5min.) and incubated with DAPI (1µg/mL, Thermo Scientific, #62248), Phalloidin-Alexa Fluor® 647 (1:100; Cell Signaling, #8940) and goat anti-rabbit Alexa Fluor® 568 (1:500; Invitrogen, #A11011; RRID: AB_143157) in 1.5 % BSA in PBS-T (1.5h, RT). After washing (3x 5min. in PBS-T), sections were mounted using Prolong Glass (Invitrogen, # P36982) and cured overnight at room temperature prior to imaging. Image acquisition was conducted in a Zeiss LSM980-Airyscan2, using 405 (DAPI), 488 (eGFP), 561 (Alexa Fluor-647) and 639nm (Alexa Fluor-647) laser lines and 63x 1.4 NA Plan-Apochromat oil immersion objective in Airyscan SR mode. Serial sections of 1.2 µm were collected with 0.21 µm step size in Zeiss’s ZEN Blue v3.0. After SR acquisition, additional tile-scan series of the whole section were obtained using the same laser lines but with 10x 0.3 NA, Plan-Neofluar objective and utilizing two PMT and one GaAsP detectors. Subsequent image analysis was performed using ImageJ (Rasband, W.S., ImageJ, U.S. NIH, Bethesda, Maryland, USA).

### Immunofluorescence *of cultured RPTEC*

RPTECs were isolated from C57BL/6J mice as described above, seeded onto µ-Slide 8 Well coverslips (Ibidi, cat# 80826), treated with heme as described above, rinsed in PBS and fixed (4% paraformaldehyde, Sigma, in PBS, 15min., RT), permeabilized (0.1% Triton X-100, Sigma, in PBS, 20min., RT), blocked with 2% goat serum (Life technologies, #16210-064) in PBS (1h, RT) and incubated with primary antibody: rabbit polyclonal anti-Slc40a1^55^ (1:250 in PBS, 2% goat serum; overnight at 4°C). Slides were washed (1x in PBS, 5min.) and incubated with secondary antibody (4µg/mL, Goat anti-Rabbit IgG (H+L) Cross-Adsorbed Secondary Antibody, Alexa Fluor 488, Invitrogen, #A11008, RRID: AB_143165) diluted in in PBS 2% goat serum (1.5h, RT). After washing with PBS (1x 5min.) slides were incubated with DAPI (0.5µg/mL, 15min.), rinsed (1x 5 min.) and mounted (MOWIOL-DABCO, MM-125). Images were captured on a Nikon Ti microscope, based on Andor Zyla 4.2 sCMOS 4.2Mpx camera controlled with the Nikon Elements software, using a 20X or 100X objective, and DAPI+FITC+Cy5 filter sets. Image analysis was performed using ImageJ (Rasband, W.S., ImageJ, U.S. NIH, Bethesda, Maryland, USA).

### Erythrocytic Compartment Preparation

Femurs were harvested from *Slc40a1^PepckΔ/Δ^* and *Slc40a1^fl/fl^* mice either not infected (NI, controls) or at 7 or 10 days after *Pcc* infection, cut and punched at both ends with a 20G needle. Bone marrow was flushed and recovered by centrifugation (2,300*g*, 10sec., RT), re-suspended in PBS (1.5mL) and passed through a cell strainer (70µm, Corning). Spleens from the same mice were harvested and grinded on a cell strainer (70µm, Corning), washed in PBS (10mL), stained and analyzed by flow cytometry (See **Flow Cytometry**).

### Tissue Leukocyte Isolation

Kidneys and livers were minced into small pieces and digested (1mL; RPMI, Gibco^TM^ ThermoFisher Scientific # 61870044, 0.2 mg/mL Liberase, Roche # 5401127001, 10 µg/mL DNase I, Roche # 10104159001) under shaking (220rpm, 37°C, 20 min). Digested tissue was passed through a cell strainer (70µm, Corning) and washed (5mL RPMI) and centrifuged (300*g*, 5min., 4°C). Pellets were washed (PBS, 0.5mM EDTA), re-suspended in 40% Percoll (6mL, GE Healthcare # 10607095; 36 vol. Percoll: 4 vol. 10X PBS: 60 vol. PBS) and carefully laid onto 80% Percoll solution (2mL, 72 vol. Percoll: 2 vol. 10X PBS: 20 vol. PBS). Samples were centrifuged (700*g*, 20min., RT, no acceleration or break). Leukocytes were collected from the ring at the interface of the 40% and 80% Percoll solutions, washed in 0.5mM EDTA PBS and stained for flow cytometry analysis (See **Flow Cytometry**). Splenocyte suspensions were obtained as mentioned above (see Erythrocytic Compartment Preparation).

### Flow Cytometry

#### Erythropoiesis panel

Cells isolated as described in **Erythrocytic Compartment Preparation** were incubated with Fc block (anti-CD16/CD32, clone 2.4G2, produced in house, 1:100) in PBS (10min., RT), followed by incubation (10 min. on ice) with fluorochrome-conjugated antibodies against: CD44 (clone 1M7, PE-conjugated, BD Pharmingen #553132, RRID: AB_394647), Ter119 (BV421-conjugated, Biolegend, #116233, RRID: AB_10933426), CD3 (clone 145.2C11, biotin-conjugated, produced in house), CD19 (clone 1D3, biotin-conjugated, BD Pharmingen #553784, RRID: AB_395048), CD11b (clone M1/70, biotin-conjugated, BD Pharmingen #553309, RRID: AB_394773), Gr1 (clone RB6.968, biotin-conjugated, BD Pharmingen #553125, RRID: AB_394641), CD49b (clone HMα2, biotin-conjugated, Biolegend, #103521, RRID: AB_2566365) and CD11c (clone N418, biotin-conjugated, Biolegend, #117303, RRID: AB_313772) at the ratio of 1:200 in PBS. Cells were washed (PBS; twice), centrifuged (300*g;* 5min.; 4°C) and stained (10min., RT) with Streptavidin (Alexa Fluor^®^ 647-conjugated, Biolegend, #405237; or PerCP-eFluor^TM^-conjugated, eBiosciences, #46-4317-82, RRID: AB_10598051) to lineage markers (1:200 in PBS). Cells were washed in PBS, centrifuged (300*g*; 5min.; 4°C) and incubated (10 min., RT) with LIVE/DEAD™ Fixable Yellow Dead Cell Stain Kit (ThermoFisher Scientific, #L34959; 1:5000 in PBS). Cell acquisition was performed in a BD LSR Fortessa X-20 (BD Biosciences) flow cytometer and data analyzed using FlowJo software V10. Erythroid cell populations in spleen and bone marrow were identified by exclusion of lineage markers (CD3, CD19, CD11b, GR1, CD49b and CD11c) and expression of Ter119 and CD44 and size, using the gating strategy illustrated in *Figure S_1*. Fe quantification in erythroblasts was performed using the FeRhoNox™-1 probe (Goryochemical, # GC901). Briefly, cells were incubated with the FeRhoNox™-1 probe (1:200 in PBS, 30min., 37°C), washed with PBS and stained for the erythropoiesis panel described above and illustrated in *Figure S_3*.

#### Hemophagocytic macrophage panel

Cells isolated as described in “Erythrocytic Compartment Preparation” and “Tissue Leukocyte Isolation” were incubated with Fc block (1:100) and LIVE/DEAD™ Fixable Yellow Dead Cell Stain Kit (ThermoFisher Scientific, #L34959; 1:5000 in PBS)(10min., RT), followed by incubation with fluorochrome-conjugated antibodies against: CD11b (FITC-conjugated, produced in house), F4/80 (clone A3-1, A647-conjugated, produced in house), Ly-6G (PE-CY7-conjugated, BD Pharmingen^TM^, #560601, RRID: AB_1727562), Ly6C (BV711-conjugated, Biolegend, #128037, RRID:AB_2562630) and CD163 (PerCP-eFluor710-conjugated, ThermoFisher Scientific, #46-1639-42, RRID:AB_2573722) at the ratio of 1:100 in PBS (in the dark, 15 min., on ice). Cells were then washed in PBS, centrifuged (300*g*, 5min. 4°C), fixed using Fix-Perm solution (100µL; eBioscience™ Foxp3 / Transcription Factor Staining Buffer Set, thermoFisher, #00-5523-00, in the dark, 35min., RT) washed in 1X Perm buffer (eBioscience™ Foxp3 / Transcription Factor Staining Buffer Set, ThermoFisher, #00-5523-00, 350*g*, 5min., RT) and incubated with anti-Ter119 (4µg/mL, BV421-conjugated, Biolegend, #116233, RRID: AB_10933426) in 1X Perm buffer (eBioscience™ Foxp3 / Transcription Factor Staining Buffer Set, ThermoFisher, #00-5523-00, in the dark, 45min., RT). Cells were washed in 1X Perm buffer (350*g*, 5min., RT), washed in PBS (350*g*, 5 min, RT) and resuspended in PBS for flow cytometry analysis. Cell acquisition was performed using a CYTEK® Aurora (Cytek Biosciences) spectral flow cytometer and data were analyzed using FlowJo software V10. Hemophagocytic macrophages in spleen and tissues were identified by exclusion of lineage markers (Ly6G) and expression of CD11b, F4/80, CD163 and Ter119, using the gating strategy illustrated in *Figure S_2*.

### Fe deficient diet

*Pcc*-infected C57Bl6 mice were fed *ad libitum* on standard chow diet *vs.* Fe-deficient diet (ssniff® EF R/M Iron deficient, ssniff Spezialdiäten GmbH, # E15510-24) starting 7 days post-*Pcc* infection. At day 10 post *Pcc* infection, mice were sacrificed, and heparinized blood was collected for hemogram analyzes, while bone marrow and spleen were collected for erythropoiesis determination by flow cytometry (see **Flow cytometry**).

### RBC supplementation experiments

Blood was collected from naïve (non-infected) C57BL6 mice and transferred (8×10^8^ RBC in 500µL PBS, i.p.) *Pcc*-infected *Slc40a1^Pepck^*^Δ/Δ^ mice, at days 6, 7 and 8-post infection. Control *Pcc*-infected *Slc40a1^Pepck^*^Δ/Δ^ and *Slc40a1^fl/fl^* mice received PBS (500µL, i.p.). Mice were monitored daily for quantification of disease parameters, pathogen load and survival, as described in ***Plasmodium* infection disease and assessment.**

### Human study design and ethical statement

This study was conducted as a cross-sectional study and quantitative approach. The study was approved by the Human Research Ethics Committee of the Institute of Health Sciences (Official Letter No. 755/GD/ISCISA/018) and authorized by the Clinician direction of Hospital Josina Machel-Maria Pia in Luanda (Official Letter No. 260/DPC/HJM/2018). This hospital is considered a national reference institution of tertiary level, receiving referrals from patients from all over the country and includes basic facilities for outpatient services, inpatient services and special services including an intensive care unit, coronary care unit, hemodynamics unit, cardiac surgery and hemodialysis unit. All patients enrolled in the study signed an informed consent on the nature and objectives of the study.

### Patient recruitment

The study population consisted of 400 of the 410 patients treated and admitted for malaria at the Josina Machel-Maria Pia Hospital between December 2018 and January 2020, including patients described^57^. Only patients who met the selection criteria and agreed to participate in the study were included. Additional information was collected through an open and closed questionnaire for patients aged 12 to 50 years old and only patients who were hospitalized for more ≥ 4 days were included. Selection criteria at enrolment included: confirmed *P. falciparum* infection, clinical symptoms including fever, headache, and other presentations of sickness behavior such as myalgia, asthenia, malaise. Patients who developed cerebral malaria or had a prior clinical history of chronic or acute kidney disease, polycystic kidney disease, hypertension, diabetes, HIV and/or other cofactors of kidney disease were excluded from further analyzes. Individuals with **sickle cell anemia** were also, excluded based on an initial screening by a sickling test and upon confirmation by gel electrophoresis. Sickle cell trait individuals were not explicitly identified and are thus included in the analyzes.

### Malaria diagnosis

*P. falciparum* infection was detected using a rapid malaria antigen test (SD-Bioline Malaria AG Pf/PAN, Abbott, # 05FK60) and confirmed microscopically in Giemsa-stained thick blood smears. Parasitemias were estimated based on the number of infected RBC detected by field (one iRBC per field corresponding to 40 iRBC/mm^3^)^57^.

### Patient follow-up and monitoring

Vital signs were assessed 3 times a day, as described^57^. Body temperature was monitored using a digital thermometer and reassessed using a mercury thermometer. Blood pressure and pulse were measured using a digital pulse sphygmomanometer and reassessed using a manual pressure sphygmomanometer or a hospital cardiac monitor, respectively. Respiratory movements were assessed by observing the chest expansion with the aid of a timer.

Hematology examinations were performed using an automatic hematology analyzer (Mindray BC-30, CBC-3-DIFF, 21 parameters and 3 histograms). Erythrogram and white blood cell data were evaluated on admission before starting antimalarial treatment. For erythrogram data, hemoglobin (Hb), red blood cell count (RBC), Red Cell Distribution Width (RDW), hematocrit count, mean corpuscular volume (MCV), mean corpuscular hemoglobin (MCH) and mean corpuscular hemoglobin concentration (MCHC) were evaluated. For the Leukogram data, lymphocyte, platelet, neutrophil and leukocyte counts were evaluated. Serum urea and creatinine were quantified in peripheral blood, using automated devices (Flexor E180 and Flexor E450, Vital Scientific), at the central laboratory at Josina Machel Hospital of Luanda, Angola.

### Acute kidney injury (AKI)

was diagnosed according to the Kidney Disease Improving Global Outcomes (KDIGO) classification system^58^. Briefly an increase in serum creatinine ≥1.5 times the expected baseline or ≥ 3,0 mg/dL, known or presumed to have occurred within the prior seven days. As preadmission creatinine values were not available, expected baseline creatinine values were calculated as recommended using the Modification of Diet in Renal Disease (MDRD) formula assuming a glomerular filtration rate (GFR) of 75 ml/min/1.73m2 for patients aging ≥ 19 years and a GFR of 100 ml/min/1.73m2 patients aging 12-18 years. **Stage 1 AKI** was defined by an increase to 1.5-1.9 times the expected baseline or an increase in creatinine to ≥3 mg/dL, **Stage 2 AKI** by an increase to 2.0–2.9 times the expected baseline and **Stage 3 AKI** by either an increase to ≥ 3 times the expected baseline or an increase in creatinine to ≥4 mg/dL^58^.

### Statistical analysis

Statistically significant differences between two groups were assessed using a two-tailed unpaired Mann-Whitney or a Welch’s t test, more than two groups using one-way ANOVA and in grouped analysis using two-way ANOVA with Sidak’s multiple comparison test. Significant differences in frequencies were determined using the Chi square test. Survival curves are represented by Kaplan– Meier plots, and the survival difference between the groups was compared using the log-rank test. All statistical analyses were performed using GraphPad Prism 7 software. Differences were considered statistically significant at a P value <0.05. NS: Not significant, P >0.05; *: P<0.05, **: P<0.01; ***: P<0.001.

### Regression analysis

To quantify the associations between urea and each of the hemogram outputs while controlling for age, gender, and parasitemia, we performed a series of multiple linear regressions with each of the hemogram outputs as response variables and urea, parasitemia, age, gender, and AKI status as independent variables. All data, except AKI status and gender, were log-transformed and standardized to allow comparison between different regressions. For each independent variable, its regression coefficient quantifies the association between that independent variable and the response variable (eg. hemoglobine.g. hem) adjusting for the remaining independent variables (age, gender, etc.).

Out of the 400 patients, 39 were excluded from the regression analysis because they were missing either age, gender or parasitemia. For each of the response variables, patients with missing information for that particular response variable were excluded. Numbers used were as follows:

| Response variable | Number patients |
| --- | --- |
| RBC | 359 |
| Hb | 361 |
| MCH | 295 |
| MCV | 358 |
| MCHC | 356 |
| RDW | 337 |

 A logistic regression was also performed to estimate probability of death using all the hemogram parameters as predictor variables and controlling for age and sex. Regressions were performed using a Bayesian framework and posterior distributions inferred through MCMC sampling. Linear regressions were performed using the python package pymc3 version 3.11.1.

## Acknowledgements

We are indebted to Dr. Joanne Thompson (University of Edinburgh) for the PcAS-GFP_ML_ parasites, Dr. Ana Domingos (Oxford University) for the *Egfp-L10^fl/wt^* mice, IGC’s Advanced Imaging; Antibody&Flow Cytometry, Genomics, Bioinformatics, Electron Microscopy, Quantitative Biology and Histopathology core facilities for excellent support. QW was supported by Marie Skłodowska-Curie Research Fellowship (RIGM 892773) and The International Postdoctoral Exchange Fellowship Program from the Peoplés Republic of China (20190090), SR by Fundação para a Ciência e Tecnologia (FCT; 5723/2014; FEDER029411), TWA by the Gulbenkian foundation (IBB2017). The MPS laboratory at Instituto Gulbenkian de Ciência is supported by the Gulbenkian, “La Caixa” (HR18-00502) and FCT (5723/2014; FEDER029411) foundations as well as by Oeiras-ERC Frontier Research Incentive Awards. MPS is an associate member of the DFG Cluster of Excellence ‘Balance of the Microverse’ (https://microverse-cluster.de/en). Support by Congento (LISBOA-01-0145-FEDER-022170) is acknowledged. Clinical data in was developed by ES team in Angola was funded by the International Society of Nephrology (ISN; Research and Prevention Program and Saving Young Lives). ES was supported by the “Envolve Science” 2021 Program of the Calouste Gulbenkian Foundation.

## Author contribution

QW: Study design, experimental work, data analyzes, interpretation and writing of the manuscript. ES: Clinical study. LVS: Experimental work, analyzes and interpretation of compensatory erythropoiesis, initial analyzes of clinical data. SC: Generation and characterization of *Slc40a1^fl/fl^* mice. TWA: Indirect calorimetry measurements, analyzes and interpretation. RM: RNAseq data analyzes and data interpretation. TP. Quantitative analyses of clinical data. JL and CN: Staining and Airyscan-image acquisition of kidney cryosection. SR: Formulation of the original hypothesis, study design, analyzes, interpretation and writing of the manuscript. PLT: Electron microscopy analyzes and interpretation. GW and FW: Conceptual input and study design. MPS: Study design, supervision of experimental work, data analyzes, interpretation and co-writing of the manuscript. All authors read and approved the manuscript.

