## Supplementary figures and images for "A Protective Inter-Organ Communication Response Against Life-Threatening Malarial Anemia"

### western blot raw data

Figure S3C RAW data

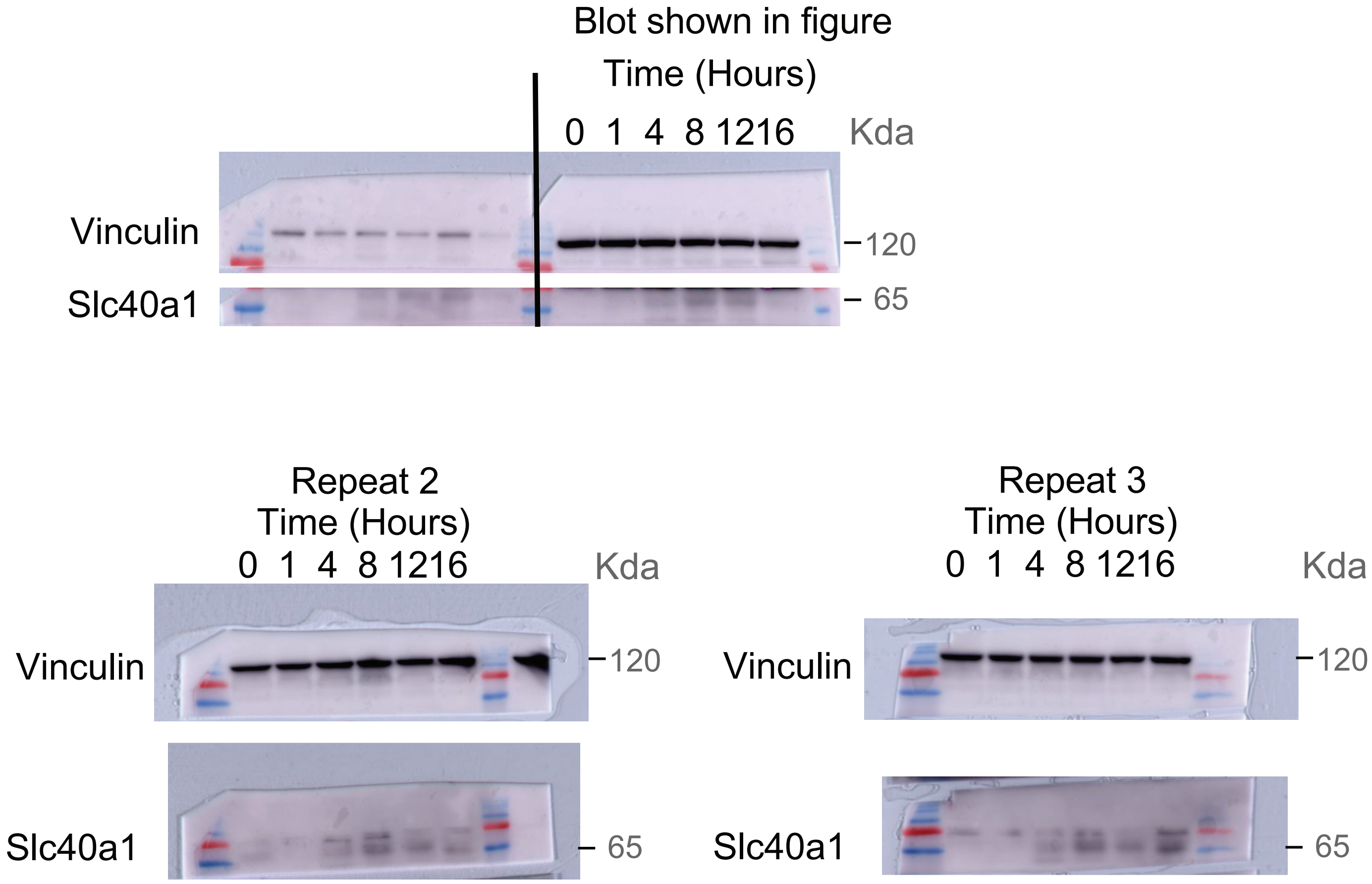


Figure S3E RAW data

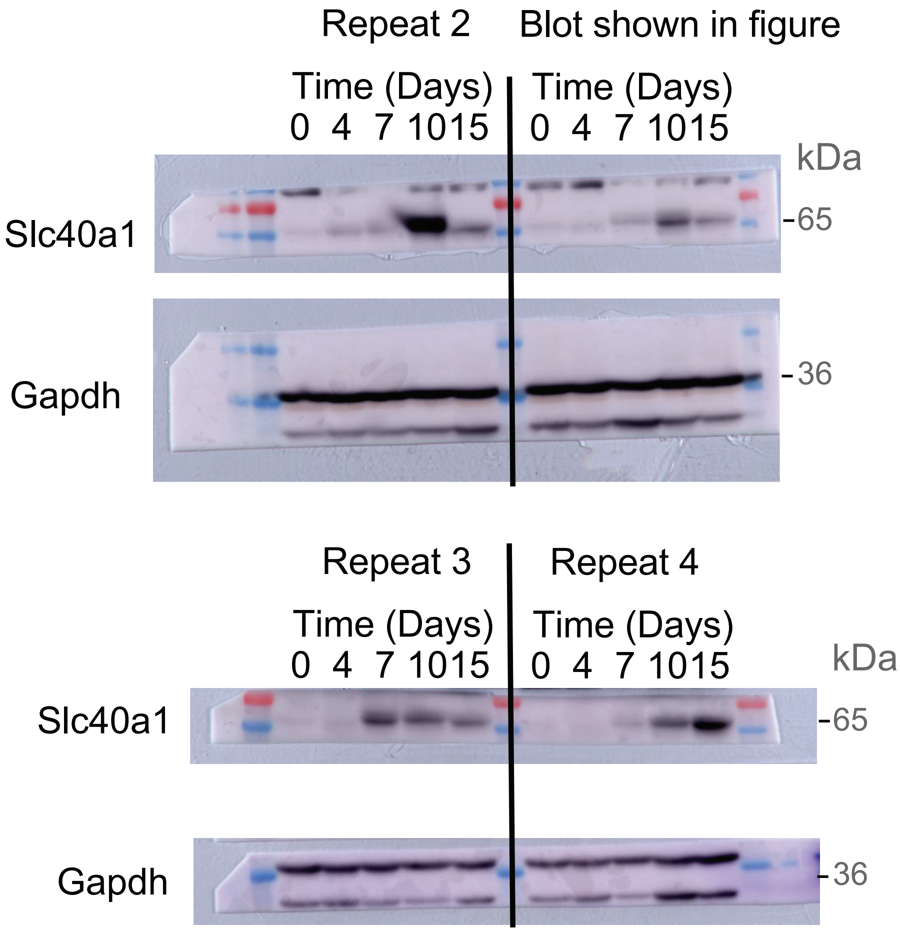
